# Prediction of transformative breakthroughs in biomedical research

**DOI:** 10.64898/2025.12.16.694385

**Authors:** Matthew T. Davis, Brad L. Busse, Salsabil Arabi, Payam Meyer, Travis A. Hoppe, Rebecca A. Meseroll, B. Ian Hutchins, Kristine A. Willis, George M. Santangelo

**Affiliations:** Office of Portfolio Analysis, Division of Program Coordination, Planning, and Strategic Initiatives, National Institutes of Health; Bethesda, MD 20892; Information School, School of Computer, Data, and Information Sciences, College of Letters and Science; University of Wisconsin-Madison, Madison, WI 53706; National Cancer Institute, National Institutes of Health; Rockville, MD 20850

**Keywords:** predictive analytics, metascience, science of science, systems science, machine learning, artificial intelligence, co-citation networks, transformative breakthroughs

## Abstract

The ability to predict scientific breakthroughs at scale would accelerate the pace of discovery and improve the efficiency of research investments. Recent advances in artificial intelligence, graph theory, and computing power have provided new ways to pursue this elusive goal. We have identified a common signature within co-citation networks that accurately predicts the occurrence of breakthroughs in medical research, on average more than 5 years in advance of the subsequent publication(s) that announced the discovery. A combination of features produces these diagnostic signals: a burst of papers exploring a novel scientific concept, an unusually high number of very influential papers in specialty journals, and low topical cohesion of the associated content. We analyzed two different periods separated by 20 years to show that the kinetics of breakthrough formation are conserved, suggesting that our approach can be used to predict which topics will produce future transformative discoveries.

**Significance statement:** Scientific breakthroughs are rare, as is contemporaneous recognition of their initial expression. Faster, more efficient identification of topics likely to produce future breakthroughs would speed scientific and technological progress. We introduce an AI/ML-detected signature in co-citation networks that recognizes such topics up to twelve years before the breakthrough itself occurs. Our findings illustrate how a better understanding of the scientific process may lead to greater scientific returns.

## Main Text

Scientific advancements occur when specific insights or ideas survive rigorous testing of their validity. Although scientists have many ideas, only a few give rise to a transformative breakthrough—that is, a small subset of scientific ideas ultimately go on to make major contributions to human health. The time, effort, and resources required to convert the germ of an idea to practice are generally recognized as rate-limiting factors in advancing scientific knowledge; an equally important but less well appreciated barrier is the delay in identifying truly transformative ideas and ensuring that they receive adequate support, especially at early stages when they are at the greatest risk of being under-appreciated.

Delays in recognizing breakthroughs in medical research are not uncommon. A prominent example was the discovery of *Helicobacter pylori* as a causal agent in chronic gastritis, peptic ulcers, and gastric adenocarcinoma. In the decade following Warren and Marshall’s Nobel prize-winning publication in 1984 (*1*), the U.S. National Institutes of Health (NIH) funded fewer than a dozen *H. pylori* research projects. Within a few years of the pair receiving their award in 2005, the agency was funding an average of a hundred such projects each year. A second example from the 1980s was delayed recognition of the need to fund prion research after that seminal idea first appeared in the literature. In the seven years following the announcement of Prusiner’s discovery in 1982 (*2*), only six proposals to study prions were funded by NIH; the current investment includes hundreds of such research awards annually. Scientists, including decision-makers at funding agencies, would therefore benefit from a systematic way to identify research topics that are (or will be) producing transformative breakthroughs.

Given the exponential growth of the PubMed database over the past six decades—over a million papers have been added each year since 2013—human curation is an increasingly inefficient way to detect trailblazing discoveries. Computational approaches (*3–8*) can now be used to supplement tasks that would otherwise require manual curation, for example in attempting to characterize a pre-specified topic that has emerged in the literature (*9, 10*). A recent report described a more ambitious attempt to identify “impactful” biotechnology research computationally (*11*), though the input variables on which that algorithm was trained have raised concerns about bias in the model’s results. In particular, the choice to limit the training data to publications from just a few dozen journals invites questions about which features are influencing its predictions (*12–16*).

The challenge of detecting new breakthroughs across the entire medical research landscape is on an entirely different scale than attempting to map breakthroughs within a defined subfield, not least because accurate mapping of those events at scale is a daunting “needle in the haystack” challenge. Transformative discoveries are rare, and the evidence of their occurrence must be mined from a very large database. Of the vast number of insights reported by scientists annually, only a few achieve “breakthrough of the year” status (*17, 18*).

To detect transformative breakthroughs at scale, we created a comprehensive network (graph) of the PubMed database, in which publications (nodes) are connected by co-citation links (edges); as expected (*19*), the weight assigned to each edge captures the degree to which the content of the two papers it connects is similar. Both subject matter experts and artificial intelligence/machine learning (AI/ML) confirmed that publications with similar content cluster together in the network and are topically distinct from those in adjacent clusters. We have developed an algorithmic approach that detects the rare signals identifying which topics will produce transformative discoveries in future years, and were surprised to find that those signals appeared up to 12 years before publication of the key papers describing the breakthrough achievement. Analysis of time frames 20 years apart (1994-1997 and 2014-2017) revealed that topics signaling future breakthroughs have shifted to more translationally focused areas of research (*20*).

### Mapping scientific topics across the entire PubMed database

The first step in detecting signals of future breakthroughs was to convert the information in scientific publications into a structured set of features amenable to high-throughput computational analysis. This conversion is typically accomplished either with natural language processing to link the content in publications, or by linking them through their lists of cited references (*21, 22*). Both approaches have utility, but content analysis may be confounded in large scale studies when investigators use different language to describe the same concept, especially across different eras (*21*). Using citation networks avoids this problem, as it has the advantage of leveraging links between publications that practicing scientists themselves create as they pursue similar or co-occurring ideas (*19, 23, 24*). Though citation networks have been used to identify features of transformative science within certain biomedical subdisciplines (*3, 19, 25–29*), doing so comprehensively across all biomedical sciences requires the processing of tens of millions of publications in the PubMed database, a challenge that is larger by several orders of magnitude. The computing power needed to address this problem at scale is now available (*30*).

Co-citation networks (CCNs) have several advantages over other types of citation networks for this task (*19, 31*). To begin, since multiple papers are added to the co-citation network of a paper with each citation it receives, CCNs are less vulnerable to distortion caused by finite number effects than networks of direct citations. Next, CCNs can also be made resistant to the distortions introduced by comparing a well-cited paper to a less-well cited one. Briefly, this is done by converting each paper to a sparse vector representing the total number of times it is cited, the identity of each paper with which it is co-cited, and the number of those co-citations, and then calculating the cosine similarity of all vectors pairwise (see Materials and Methods and Supplementary Text for details). Finally, CCNs are highly dynamic; as new discoveries connect the content of a paper to neighboring areas of investigation, that paper gains additional citations and becomes increasingly influential. As a paper becomes more influential, it in turn draws more attention from scientists working to build on established knowledge in a wider range of disciplines. In this way, CCNs robustly capture the intellectual content of an influential paper, both as it was originally conceived by the authors and as it is assimilated, interpreted, and re-interpreted by one, several, or even many interested scholarly sub-communities as science progresses and knowledge about the corresponding topic improves.

We therefore began by constructing a CCN of the 17.2M cited papers in PubMed for all years, beginning with the oldest publications in the database and running through the end of 2017. Due to the nature of CCNs, publications on the same topic form a subnetwork or cluster with high edge density and short path lengths. Clusters that represent closely related but distinct topics are connected by edges with high betweenness centrality (i.e., the edges representing the shortest path between two nodes, which is the most robust and commonly accepted measure of centrality in a network (*32*); Fig. 1A). After testing several methods (Supplemental Text, fig. S1-S4), we chose a low inflation setting of the Regularized Markov Cluster (RMCL) algorithm (*33–35*) as the best means of identifying the high betweenness edges that bridge distinct topics. These edges agnostically and definitively establish the borders between topics, as opposed to clustering methods like k-means that require the number of topics to be (often arbitrarily) pre-defined. RMCL analysis indicated that the 17.2M papers in our network represent a total of 23,867 connected topic clusters, each of which is linked directly to a median of 5 (mean of 8.9) nearest neighbor topic clusters.

**Fig. 1.**
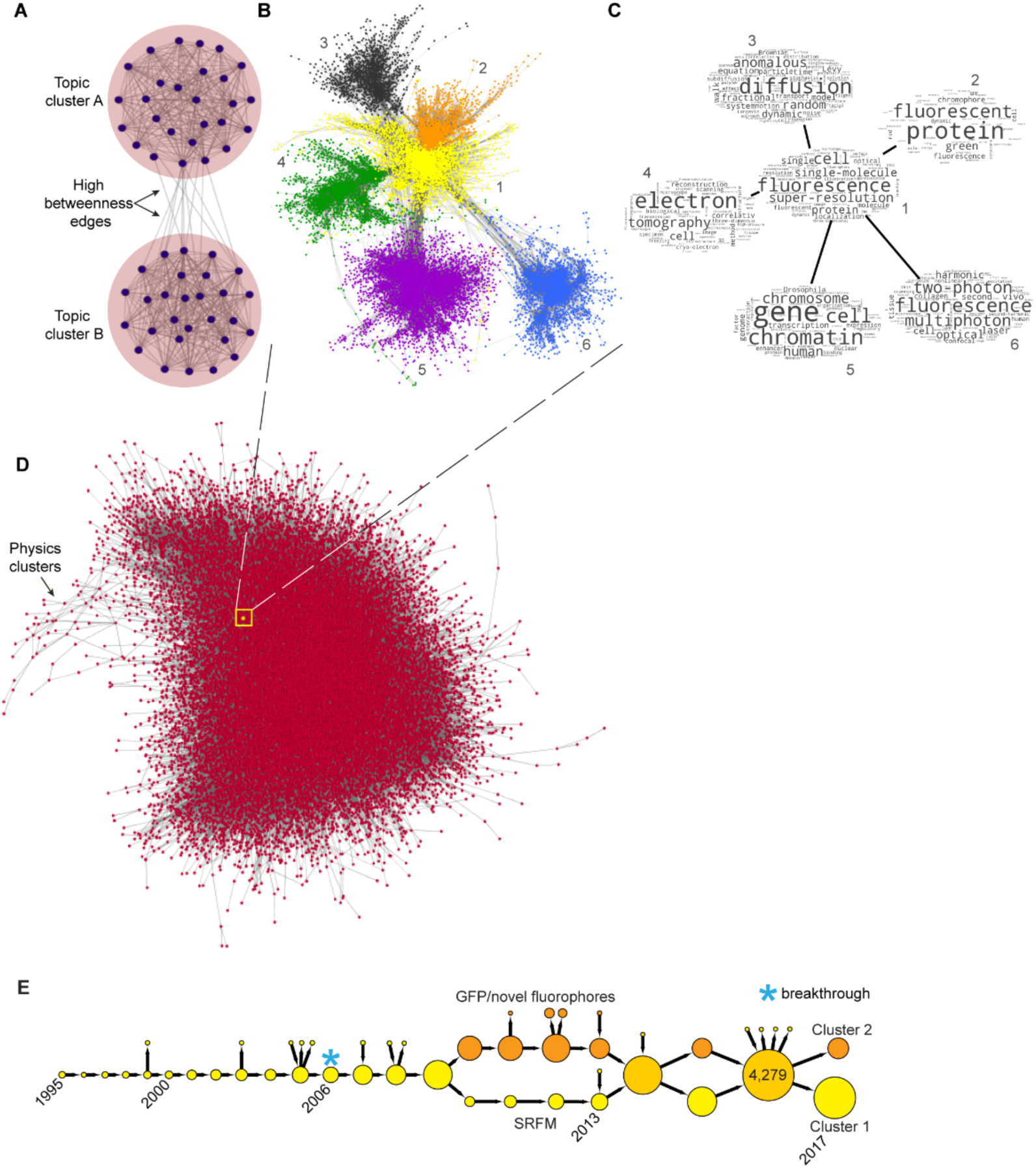
Topic clusters in the PubMed co-citation network (CCN). **(A)** Publications (small black dots) assigned to the same CCN topic cluster (large pink circles) by RMCL (see Materials and Methods) have a high density of topical links (black lines) that result in short path lengths; closely related nearest neighbors, such as topic clusters A and B shown here, are connected by high-betweenness edges; **(B)** The complete subnetwork of the super-resolution fluorescence microscopy (SRFM) cluster (1, yellow) and five of its nearest neighbors (2 through 6, orange, black, green, purple, blue, respectively). **(C)** Clusters 1 through 6 in **(B)**, represented as word clouds of the titles and abstracts of publications in each cluster; **(D)** Coarsened version of the entire PubMed CCN/RMCL network; each red dot is analogous to the pink circles in **(A)** and represents a topic cluster with its associated intra-cluster edges, collapsed into one node; each group of high-betweenness edges that bridges two clusters is also collapsed into one edge (see Materials and Methods). The region representing sparsely covered non-biomedical topics in PubMed (e.g. particle physics and astrophysics), and the location of the SRFM cluster, are indicated by a black arrow and small yellow dot, respectively; **(E)** The SRFM/GFP trajectory from 1995 (the first year it existed as a discrete cluster) through 2017 (see Materials and Methods); the area of each cluster (circle) is proportional to the number of publications it contains, benchmarked to the size of the largest (2016) cluster of 4,279 publications. Blue asterisk indicates the cluster in which the breakthrough papers (hereafter, breakthrough cluster), which were published in 2006, first appear.

Although it is impractical to depict a network that large in its entirety, the complete network for 6 of the ∼24K linked topic clusters is shown in Fig. 1B. Those 6 clusters together represent 15,675 papers and 501,877 connecting edges, comprising 0.092% and 0.126% of the entire network, respectively. Nodes of the same color indicate papers identified by RMCL as covering the same topic; word clouds capture the themes of each of the six clusters (Fig. 1C). Note that cluster 1 (yellow subnetwork in Fig. 1B) includes publications describing the 2006 breakthrough in super-resolution (sub-diffraction limit) fluorescence microscopy (SRFM), which merited the Nobel Prize in Chemistry in 2014. A coarsened version of the entire network, including all 23,867 linked clusters, is shown in Fig. 1D.

The portion of the network displayed in Fig. 1B,C suggested that RMCL accurately assigns publications to the same CCN cluster when their semantic content is closely aligned (*27, 36*), and assigns papers on different topics to separate clusters, even if those topics are closely related. We tested this premise by analyzing a small group of disciplinary journals that focus on what are commonly accepted as discrete areas of research, and then expanded that approach to test the entire corpus. We began with 100,000 randomly selected pairs of articles from each of three disciplinary journals (Genetics, Journal of Neuroscience, and Blood), and as controls, from each of three multidisciplinary journals (Proc. Natl. Acad. Sci. [PNAS], Science, and Nature). We then used a well-established language model (*36, 37*) to calculate the semantic similarity of the two papers in each journal-matched pair.

As expected, pairs of papers selected at random from the same disciplinary journal tend to be more similar to each other than pairs selected from the same multidisciplinary journal (blue bars, fig. S5A). Since each paper was assigned to a cluster by RMCL, we could also test how the similarity of its “journal-mates” compares with that of its “cluster-mates.” Strikingly, cluster-mates are as a class more semantically similar than journal-mates (Fig. 2, fig. S5A), and as the semantic similarity between a pair of journal-mates increases, the probability that RMCL assigned them to the same cluster increases exponentially (fig. S5B). We also compared journal-mates assigned to different clusters with those that share the same cluster. Since the latter have significantly higher average similarity (Fig. 2B; light blue vs. dark blue bars, p<0.001), for the journals tested our approach separates even closely related topics. To confirm that these results extend to the full corpus, we randomly selected 599,169 pairs of papers from all RMCL clusters without any reference to journal of publication. The distribution of similarities for these randomly drawn cluster-mates approaches those for pairs that share both a cluster and a journal (Fig. 2A, compare light green to dark green curves), and far exceeds that of journal-mates (Fig. 2A, compare light green curve to curves in blue and orange tones).

**Fig. 2.**
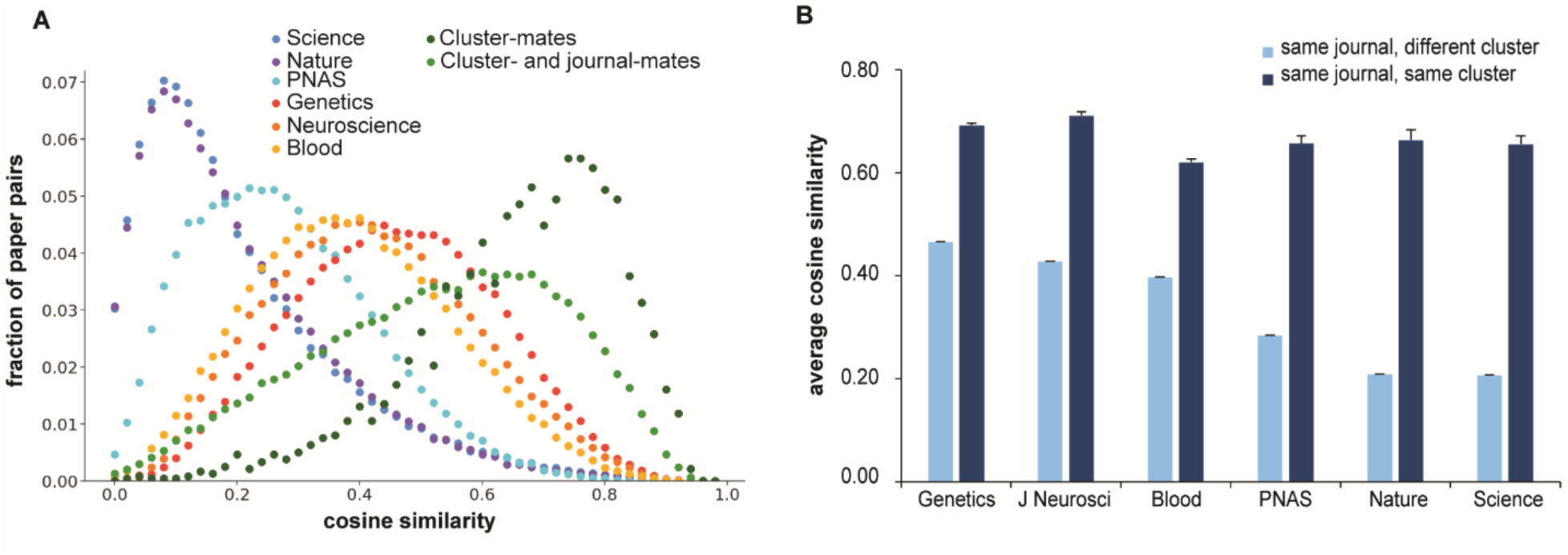
Papers in an RMCL-assigned cluster are semantically more closely related than those in the same disciplinary journal. Cosine similarity scores were calculated based on vectors generated by a well-established language model (32,33). **(A)** Distribution of cosine similarity scores. Curves shifted to the right display greater semantic similarity. Categories in descending order of semantic similarity (right to left) are as follows: papers in the same RMCL cluster (two green curves), the same disciplinary journal (Genetics, Neuroscience, Blood; three orange curves), and the same multidisciplinary journals (Science, Nature, PNAS; three blue curves). **(B)** Average cosine similarity scores for the journals listed in **(A)**. Light blue bars, papers published in the same journal but assigned to different RMCL clusters; dark blue bars, papers published in the same journal and assigned to the same RMCL cluster. Error bars, standard error of the mean (SEM).

As a final test, we provided subject matter experts with sets of papers of increasing topical similarity, and asked them to divide each set into two groups; their assignments agreed with the RMCL cluster designations, even when the publications are close to each other in multidimensional space (*36*) (Data S1). Taken together, these results demonstrate that RMCL mapping of topic clusters across the CCN network assigns papers on the same topic to the same cluster, separates even closely related topics, and organizes topics across the entire network based on the extent of their content similarity.

### Mapping past transformative breakthroughs

To determine whether topics that have already produced breakthroughs might share features that can be detected computationally, we next needed to be able to track the development of each topic from its inception through to the present year. Given the goal of identifying diagnostic signals of current or future transformative discoveries in real time, it was critical to ensure that the training set of past examples included only data that was available contemporaneously. In other words, we needed to detect past examples of diagnostic signals without relying upon information “leaked” from the future. We therefore built 36 additional historical CCN/RMCL networks, one for each year between 1981 and 2017, excluding all papers and citations beyond the terminal year (Data S2).

Once formed, individual clusters are typically quite stable; the probability that semantically similar papers assigned to the same cluster will remain together for more than thirty years is approximately 80% (Supplemental Text, fig. S6A). We therefore adopted the most straightforward means of matching each cluster to its closest counterpart in these historical networks—asking if they shared a simple majority of their papers. If so, we connected them in series, so that each cluster from the network of year x is linked to its counterparts in year x-1 and x+1 (see Materials and Methods and fig. S7 for a full description of this algorithm). For example, each 2017 cluster was linked to its closest counterpart(s) in 2016, the latter to its closest counterpart(s) in 2015, and so on back to the earliest year that the topic existed as a discrete cluster, using the entire PubMed database to create a summative chronological trajectory for each topic over that 36-year span.

The SRFM trajectory, which includes clusters 1 and 2 in Fig. 1B,C, is shown in Fig. 1E; it illustrates the rapid growth of that topic leading up to the discovery of SRFM, and includes more than 275 articles authored by the three Nobelists awarded the 2014 Chemistry prize for that transformative breakthrough (Data S3). The earliest cluster in the trajectory contains a small group of papers describing the foundational work on the use of fluorescent proteins as markers in a wide variety of biological systems. Related papers continued to accumulate until the publication of the 2006 studies that subject matter experts identify as representing the SRFM breakthrough (light blue asterisk in Fig. 1E, Data S3). The SRFM trajectory split into two branches in 2010 as the ongoing engineering of novel fluorophores diverged from the related development of new fluorescent tracking methodologies, including analysis at the level of single cells or single molecules. The repeated mergers and splits between 2014 and 2017 are an indication that these two closely related topics remained distinct but intertwined.

While the above example suggests that our trajectories faithfully represent the historical development of each topic in the PubMed database, we wanted to verify that this was broadly true before going further. One of the many advantageous features of the PubMed database is that, since 1961, the National Library of Medicine has flagged the subset of publications that are review papers. Taking advantage of those structured data, we used our language model (*36, 37*) to analyze the content of review papers in each trajectory; this leveraged at scale the subject matter expertise of scientists, who over the course of the trajectory have published those curated summaries of the literature in their respective areas of research. The expectation *a priori* was that the content of review papers would align most closely with the content of the research articles present in the same trajectory, even when comparing trajectories that represent the most closely related but distinct topics (i.e., nearest neighbors in multidimensional space). The results, which were done reciprocally for 100 randomly selected trajectories and their nearest neighbors, confirmed this expectation (fig. S8). The CCN/RMCL-based trajectories therefore accurately distinguish the development of even closely related topics over time as new reports appear in the literature.

We next identified trajectories that represented 21 different examples of biomedically relevant topics that produced breakthroughs, including the SRFM example shown in Figure 1E. Each of these achievements, which together spanned a 30-year time frame (1984 through 2014), was associated with one or more prestigious awards (13 received Nobel prizes and an additional 3 merited Lasker awards; table S1). Based upon the descriptions accompanying the announcement of a Nobel prize, Lasker award, or other formal recognition of each of these breakthroughs, we manually curated the key transformative papers and years in which they were published. This in turn determined the “breakthrough cluster” and corresponding “gold standard” trajectory that captured progress in understanding that topic, both before and after the breakthrough occurred (table S1). A sample of gold standard trajectories are shown in Fig. 1E and Fig. 3A-D (for the remaining 16, see fig. S9): the aforementioned SRFM breakthrough, exosome transfer of microRNAs (2015 ISEV Special Achievement award), cancer immunotherapy (2018 Nobel prize in Physiology or Medicine), gene editing/CRISPR (Clustered Regularly Interspaced Short Palindromic Repeats; 2020 Nobel prize in Chemistry), and drug development targeting the Epidermal Growth Factor Receptor (EGFR) oncogene (2019 Lasker award). A light blue asterisk indicates the year of those breakthrough clusters in 2006, 2008, 1996, 2014, and 2002, respectively.

**Fig. 3.**
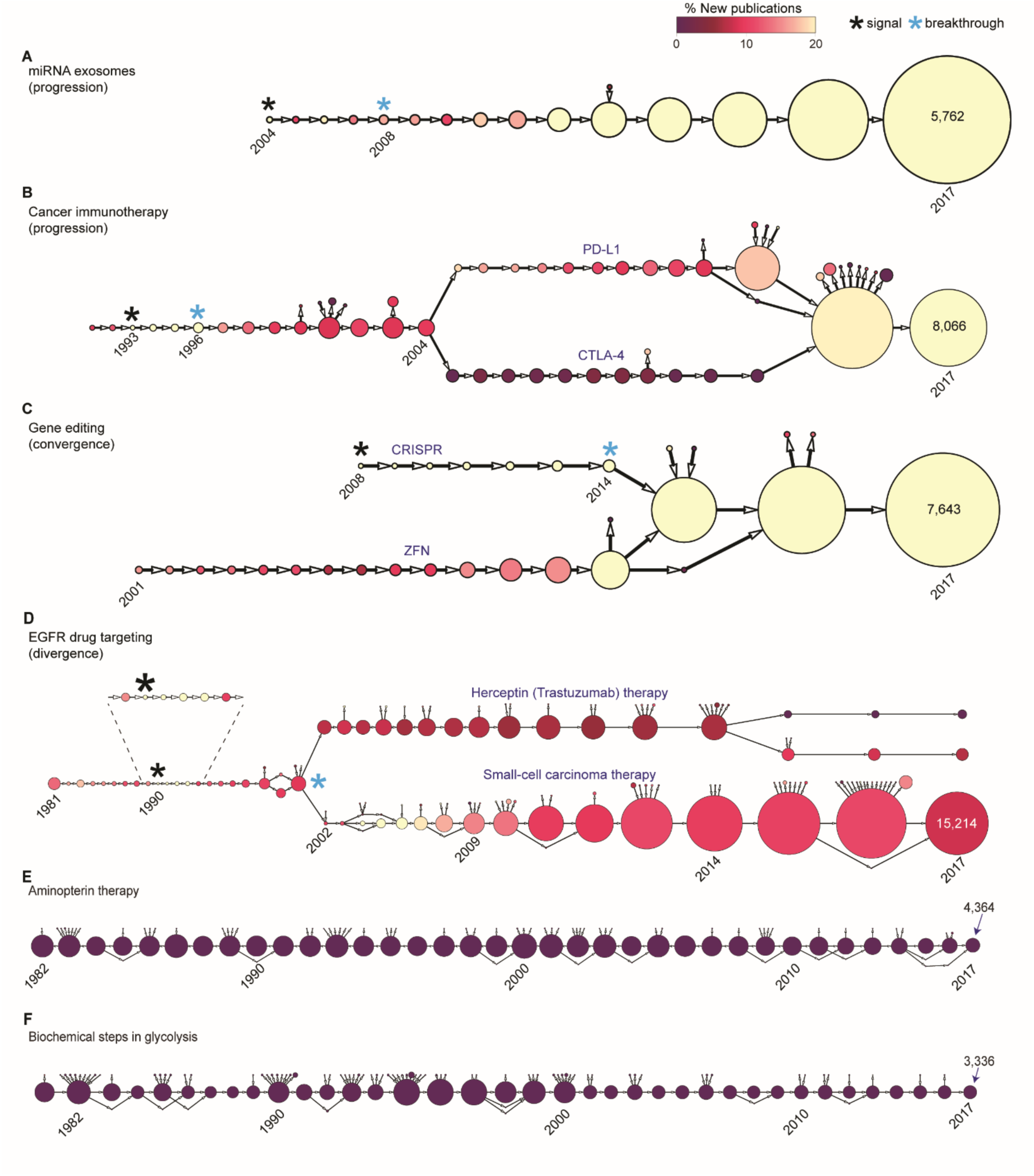
Trajectories of research topics that produced award-winning breakthroughs in biomedical research. **(A)** Exosome-mediated transfer of microRNAs; **(B)** Cancer immunotherapy; **(C)** Gene editing; **(D)** epidermal growth factor receptor (EGFR) drug targeting. Trajectories representing **(E)** Aminopterin therapy; and **(F)** Biochemical characterization of the steps in glycolysis—topics that are widely understood by practicing biomedical scientists to represent obsolete areas of research and solved science, respectively—are shown for comparison. The area of each cluster (circle) is proportional to the number of publications it contains, relative to the indicated number of publications in the largest cluster in that same trajectory. Heat mapping, which is applied uniformly to all clusters, indicates the fraction of new publications in each cluster, i.e., those that were published either in the terminal year of the cluster (x) or in year x-1; lighter colors indicate a higher fraction of new publications. Black asterisks mark clusters that display the signal of a future breakthrough, and light blue asterisks mark the subsequent cluster in each trajectory that contains the earliest paper(s) announcing the transformative discovery. The inset in **(D)** (dashed lines) is a magnified view of the trajectory in the years surrounding that breakthrough signal. The three types of trajectory—progression, convergence, and divergence (see text)—are indicated inside parentheses. Labels indicate the nature of the separate topics in years that display two major clusters in a given trajectory: T cell receptors PD-L1 vs. CTLA-4 in cancer immunotherapy; zinc finger nuclease (ZFN) vs. CRISPR approaches in gene editing, and breast cancer/trastuzumab vs. small cell carcinoma/gefitinib therapy in EGFR drug targeting. For each trajectory, the number of papers in the 2017 cluster is indicated.

Each of the 21 gold standard trajectories can be assigned to one of three types based on the nature of the breakthrough cluster and the configuration of its adjacent edges in the corresponding trajectory: progression, where each breakthrough cluster links to a single cluster in the following year (e.g. the SRFM, miRNA exosomes, and cancer immunotherapy breakthroughs in Fig. 1E, 3A, and 3B, respectively); convergence, where the breakthrough cluster merges with one or more other clusters, creating a single combined cluster (e.g. the gene editing/CRISPR breakthrough in Fig. 3C); and divergence, where the breakthrough cluster forms by splitting away from one or more other clusters (e.g. the EGFR drug targeting breakthrough in Fig. 3D). For our 21 gold standards, the distribution of the three types of breakthrough is as follows: 11 progression, 6 convergence, and 4 divergence (Fig. 1E, Fig. 3, fig. S9, and table S1). Note that we are flagging a convergence or divergence event only when the breakthrough cluster participates in the merge event or results from the split event, respectively; the cancer immunotherapy trajectory is in the progression category because the 2005 PD-L1/CTLA-4 split occurred 9 years after the breakthrough was first reported in 1996 (*38*) (Fig. 3B). Across our entire corpus, over 97% of edges are between two clusters that do not participate in a branch event and would therefore place a trajectory in the progression category if located adjacent to a breakthrough cluster. The frequency of convergence or divergence events preceding or following a breakthrough is therefore unusually high relative to the respective Bayesian priors of 0.90% (P<0.001) and 1.47% (P<0.001; see Materials and Methods).

We have defined convergence and divergence as a special case of the splitting and merging of clusters; however, splitting and merging can occur at any point over the length of a trajectory. We therefore asked whether the frequency of splits and merges could be linked to trajectory size. Controlling for the number of edges, the likelihood of branching over the entire length of our gold standard trajectories is not detectably different than the likelihood of branching in a randomly selected set of trajectories of comparable length (p = 0.08). Although not conclusive due to the relatively small number of gold standards in our sample, our data are consistent with a model where branching is not unique to breakthroughs, but breakthroughs are more likely to arise when two distinct areas of research merge or when one area divides into two or more new subfields.

### Features that signal future transformative breakthroughs

We next applied logistic regression to our gold standard trajectories to determine whether the years leading up to the breakthrough event in each topic share any commonalities that could serve as a predictive signal. An advantage of logistic regression is that, unlike neural nets or other deep learning methods, it is simple to determine the extent to which each feature contributes to the model outputs by examining its contribution to class separation (see Materials and Methods). The feature that provided the strongest signal of a coming breakthrough is a sudden upsurge in the number of publications focused on that topic (fig. S10A). This feature, which we define as %New, is quantifiable because any given cluster contains the full spectrum of papers on the topic it represents, regardless of the year in which they were published. It is therefore simple to detect, relative to historical precedent, a rapid expansion in the number of papers published in the terminal year of an individual cluster. For example, the 1996 cluster signaling the impending 2006 breakthrough in SRFM (Fig. 1E; table S1) contains papers relevant to that topic going back to 1974, so the number of papers published in calendar years 1995 and 1996 can easily be calculated as a percentage of all papers assigned to that 1996 cluster. The second important feature, which likewise captured rapidly expanding, leading-edge exploration of the topic, measures the growth of the literature contained in a cluster’s most recent ancestor, as described by its trajectory (fig. S10D). The third feature is a measure of topic cohesion (fig. S10G; calculated as entropy, see Materials and Methods). The fourth feature captures the occurrence of highly influential publications, as measured by a modified version of the Relative Citation Ratio (RCR), an article-level, field- and time-normalized indicator of scholarly recognition (fig. S10J)(*31, 39*). RCR has no citation window; it incorporates every citation received by both the article of interest and the group of peer papers to which that article is compared, making it uniquely suited for evaluating how scholarly influence evolves over time. In making predictions, however, as noted above, it is critical to avoid leaking information from the future. We therefore calculated the influence of each paper as a series of snapshots in time, year by year, using only information that would have been available contemporaneously (see Materials and Methods for details). This generates approximately 334 million historical RCR (hRCR) values for the 28 million papers we analyzed. We exclusively used hRCRs in our regression and all related analyses. These four variables together—%New, ancestral %New, topic cohesion, and influence—offer the optimal combination of predictive and explanatory power (see Materials and Methods). The mean and standard deviation of each were included in the regression as a way to incorporate temporal trends in the five years preceding each cluster, as identified by its trajectory; however, these had a minimal effect on the results (fig. S10B-C, E-F, H-I, and K-L).

Heat mapping the clusters in gold standard trajectories with %New, the strongest component of the diagnostic signal (Fig. 3A-D; fig. S9), illustrates the success of this approach in anticipating future breakthroughs in biomedical research. The cluster with the highest %New value in each trajectory in Fig. 3A-D is marked with a black asterisk; those “signal clusters” precede publication of the key breakthrough papers by up to 12 years (5.6 years on average; fig. S9 and table S1). More than a third of clusters have a %New value of 0.0; these inactive topics primarily fall into two distinct categories: obsolete, unproductive lines of investigation, or completed work that has long been accepted as a solved problem. Trajectories composed entirely of such clusters can be considered stagnant; two examples, the outdated clinical use of aminopterin as a chemotherapeutic agent and the long-ago completed characterization of biochemical steps in the glycolysis pathway, are shown as controls (Fig. 3E, F). The maintenance of high %New values across a trajectory (e.g. Fig. 3A and 3C) is particularly striking given that a larger and larger number of new papers must be added to maintain a high %New value (*40*) as the total number of papers increases over time.

All components of the breakthrough signal leverage article-, not journal-level information. Since article-level structured data have long been available, it is puzzling that journal-level metrics continue to be used despite their well-known limitations (*41–43*). One recent study with the similar goal of predicting future impact limited the set of articles it considered to have transformative potential to those published in 42 journals, selected in part based on their impact factor, a metric known to display both geographic and demographic skew (*13–16, 44*). We therefore investigated how the use of a journal-based approach might impair our ability to identify transformative research. Though it is difficult to reconstruct journal impact factors, since their calculation is not transparent (*41*), journal citation rates (JCRs) are a close approximation (*31*). We calculated the average JCR of each cluster that signaled a future breakthrough in our gold standard trajectories. Within a distribution that include 4,530 randomly selected, size-matched controls, the average JCR of the gold standard clusters ranked between the 79^th^ and the 97^th^ percentile, which means that 135 (∼3%) of those control clusters had an average JCR higher than the highest of the 21 gold standard clusters. Putting other methodological considerations aside, selecting candidates in part based on JCR would therefore have increased the number of potential false positives in our setB time frame by almost seven-fold (i.e., screening ∼160,000 clusters). In other words, basing our approach on journal-level metrics would generate enough noise to make the signal of a future breakthrough difficult or impossible to detect.

### Detection of transformative breakthroughs at scale with logistic regression

We next used logistic regression to test the possibility that, taken together, the four features we described above constitute an empirically useful diagnostic signal that can predict the occurrence of future transformative breakthroughs. We generated a model consolidating those four features into one (see Materials and Methods), and then applied that model to all 186,449 clusters in two distinct 4-year windows that are 20 years apart (setA: 1994 through 1997 and setB: 2014 through 2017). To avoid trivial or non-biomedical positives in the outputs, we limited the analysis to clusters with 40 or more publications, at least 5% of which were linked to at least one NIH award. We identified 18 and 19 clusters in setA and setB, respectively, that matched the signature of an impending breakthrough in the gold standard trajectories. This represents an average of 4.6 (median of 5) signals of future biomedical breakthroughs per year (Fig. 4A,B; table S2; see setA trajectories in fig. S11). As mentioned above for the gold standard trajectories, disciplinary journals were the most common venue of publication in signal clusters from setA and setB (82 and 94%, respectively). Values for each of the four individual features for setA and set B signal clusters, as well as for gold standard clusters and a matched sample of negative controls, are listed in Data S4; the corresponding distributions are compared in figure S10. To visualize the relative positions of these newly identified signal clusters within the biomedical landscape, we plotted the strongest component of the integrated signal, the percentage of new publications (%New), versus the prevalence of Human Medical Subject Heading (MeSH) terms (Fig. 4A,B) and translational potential of the corresponding publications (the latter defined by mean Approximate Potential to Translate, or APT, score; Fig. 4C,D). As a rule, clusters with fewer human MeSH terms and lower APT scores represent more fundamental research topics, while more human MeSH terms and higher APT scores represent more clinically oriented topics.

**Fig. 4.**
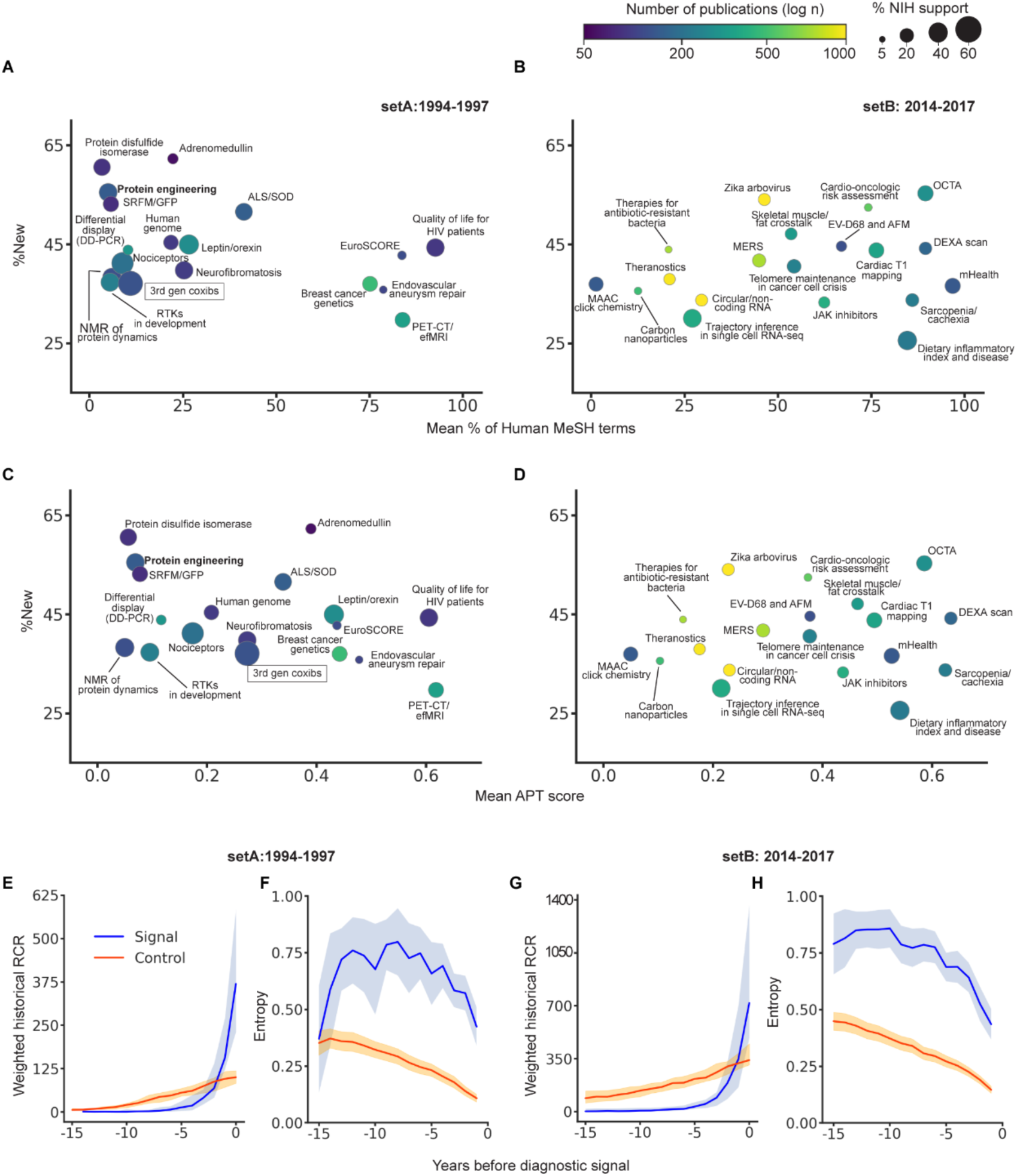
Characteristics of topics that signal future breakthroughs. The two groups of clusters matching our diagnostic signal of a coming breakthrough were setA (1994-1997; **(A,C)**) and setB, (2014-2017; **(B,D)**); the percentage of new publications (%New) is plotted against two different measures of clinical relevance (see text): the mean percentage of Human MeSH terms (**A,B**), or APT scores (**C,D**). The number of publications in each cluster, which is the denominator in our calculation of %New, is indicated by the rainbow heat map. Cluster size indicates the percentage of papers that report NIH support for the work. The weighted historical RCR (**E,G**) and entropy (**F,H**) of setA (**E,F**) and setB (**G,H**) clusters (blue lines; 95% confidence interval in light blue) are plotted for 15 years prior to the appearance of the diagnostic signal. 370 clusters containing the same number of papers, +/- .01%, as the 37 signal clusters are shown as a size-matched but otherwise randomly selected control (orange lines; 95% confidence intervals in light orange). The approach used to generate the data labels in (A-D) is described in Materials and Methods.

Of the 18 diagnostic signals identified in the setA time frame, we previously identified three as gold standards: breast cancer genetics (1994 signal; 1995 breakthrough; 2014 Lasker award; fig. S9h), leptin/orexin (1996 signal; 1998 breakthrough; 2010 Lasker award; fig. S9l), and SRFM (1995 signal; 2006 breakthrough; 2014 Nobel prize in Chemistry; Fig. 1E). Of the remaining 15, 11 have received prizes or awards and could have been included in our training set if we had encountered them initially, including the Nobel prize in Chemistry in 2018, which recognized three scientists for their groundbreaking work on protein engineering (Fig. 4, bold text; table S2). Two topics, receptor activity-modifying proteins (RAMPs) and Neurofibromatosis type 2, were in a group that produced translational breakthroughs leading to the development of therapeutic candidates now in clinical trials (fig. S12). Of the two remaining clusters in setA, one appears to be a false positive signal (boxed label in Fig. 4A,C) that arose as a result of the then-recent development of the coxibs Vioxx and Celebrex, non-steroidal anti-inflammatory drugs (NSAIDs) that directly target cycloxygenase-2 (COX-2); subject matter experts now agree the anticipation at the time of so-called third generation coxibs with improved effectiveness and safety profiles was at best premature, since their development has yet to come to fruition (*45*). The final setA topic is the development of tools that use quality of life assessments to improve clinical care for HIV patients.

We next asked how many other major discoveries recognized with prizes might be associated with a breakthrough signal that appeared outside of the setA and setB time frames. We compiled a comprehensive list of the Lasker awards and Nobel prizes in Medicine since 2014 that we neither used as gold standards nor discovered in setA or setB (six each; see Materials and Methods), and asked whether we would have been able to predict them in advance. The foundational breakthrough associated with the 2014 Nobel for the discovery of the neurological system that encodes spatial positioning occurred in the 1970s (*46, 47*), putting it outside the chronological range of our current dataset. Of the remaining eleven, all corresponded to a positive logistic regression signal (∼0.5% of all clusters); nine met all the criteria established by our gold standards, while two (the unfolded protein response, 2014 Lasker and aDNA, 2022 Nobel in Physiology or Medicine) were smaller and/or slower growing (table S3). Since not all discoveries that significantly advance human health are recognized by a major award, it is impossible to define the comprehensive set of true instances of breakthroughs, and therefore difficult to assess precision and recall. Nevertheless, our approach not only seems effective in identifying topics destined to receive high-profile recognition, but may surface topics that are otherwise overlooked by prize committees, e.g., quality of life care for HIV+ patients, or the use of mobile phones for healthcare delivery (in setA and setB, respectively).

### Kinetics of signal formation

After having validated that the setA signals were successful in agnostically predicting which topics would produce scientific breakthroughs, we wondered whether they all arose with the same kinetics. Since the signals were all detected with the same four diagnostic features, their formation could have followed a common pattern. Alternatively, given the variation in the size of the signal clusters (between 61 and 474 publications; table S2), the distribution across three different types of transformative events (progression, convergence, divergence), and the variable position of each breakthrough-signaling topic along the fundamental-to-clinical axis (based on the percentage of Human MeSH terms and APT scores (*20*); Fig. 4A,C), the 15 breakthroughs might have arisen in a non-uniform fashion. Another unknown aspect of signal formation was the timing of its appearance: it might arise slowly in a gradual accumulative process, or rapidly in just a few years with a burst of scholarly activity, as suggested by two of the four diagnostic variables.

We tracked the development of the signals in setA topic clusters for 15 years prior to their appearance, measuring the diagnostic features in those clusters as well as a large group of year-and size-matched controls (Fig. 4E,F; fig. S13). Weighted historical RCR (whRCR, the sum of hRCRs), which reflects both the number of investigators focusing on the topic and the overall level of interest in the resulting papers (*31*), remained below the level of control clusters until a rapid burst that began a few years prior to the appearance of the signal in year 0 (Fig. 4E; P<0.001). Topics displaying a diagnostic signal had lower topical coherence than the controls, measured as a higher level of entropy (see Materials and Methods; Fig. 4F; P<0.001), although this gap narrows over time following the appearance of the signal. Taken together, these results suggest that the signals appear with similar kinetics as the explanatory potential of each nascent research area of unusually high topical diversity is realized and culminates in a burst of papers and overall influence (whRCR).

Although it is very rare for any paper to experience a large increase in RCR over time (*31, 39*), the setA clusters include subsets of papers that exhibited unusually rapid growth in influence after year 0, eventually reaching >3-fold higher RCR levels; this is also a common feature of the clusters that signaled a breakthrough in the 21 gold standard trajectories (Data S5 and S6; P<0.001). Most of this increase in RCR occurred in the time interval between the signal and the corresponding breakthrough, suggesting that growth in the influence of key papers is a lagging echo of the diagnostic signal. A striking example of this phenomenon is the cluster that signaled a breakthrough in the topic focused on signal transduction by the peptide hormone adrenomedullin, which works through one of two receptor activity-modifying proteins (RAMPs; Fig. 4A,C). The 1993 *Biochemical and Biophysical Research Communications* paper reporting the discovery of adrenomedullin (*48*) increased in hRCR from 1.5 in 1993 to 19.0 in 1995, which likely contributed to the appearance of the diagnostic signal in the corresponding 1995 cluster (see the corresponding trajectory in fig. S11g; the RCR of that paper in November 2024 was 54.6). The subsequent breakthrough was announced in a 1998 *Nature* paper reporting the discovery of the adrenomedullin receptor and a new family of single transmembrane domain proteins called RAMPs (*49*); both adrenomedullin and RAMPs remain active areas of research, and the seminal 1993 paper continues, 27 years later, to add to its cumulative total of over 1400 citations.

Like setA, all of the diagnostic features of the 19 setB clusters (2014 through 2017) increased sharply immediately preceding the appearance of the diagnostic signal (fig. S13; Fig. 4B,D; and Fig. 4G,H). SetB clusters were also distributed across the entire fundamental-to-clinical axis, though interestingly the topics displaying diagnostic signals were overall more clinically focused than those in setA (compare B,D with A,C in Fig. 4). Given the similar properties of the two sets (Fig. 4E-H; fig. S13), we expect that many if not most of the setB signals will produce transformative breakthroughs in the corresponding topics going forward; five have already done so. Two of the signals in setB are associated with the development of new drugs – Janus kinase inhibitors, which the FDA has approved to treat several immune disorders (JAK inhibitors, Fig. 4B,C) and the recently-announced novel class of antibiotics targeting lipopolysaccharide transport in multi-drug resistant Gram negative *Enterobacteriaceae* (*50, 51*); Fig. 4B,C). A third signal corresponds to the development of a fragility score for the clinical management of osteoporosis; new software combining the World Health Organization’s Fracture Risk Assessment Tool with t- and z-scores derived from DEXA scans of the lumbar spine was approved by the FDA in 2016 (DEXA scan/trabecular bone score, Fig. 4B,C). Fourth, optical coherence tomography angiography (OCTA), a novel technology to rapidly diagnose retinal diseases, won the 2023 Lasker-DeBakey Clinical Medical Research Award (OCTA, Fig. 4B,C). The final example is a setB cluster describing the novel inclusion of Ru(II) and other metal catalysts in click chemistry, which was recognized by the 2022 Nobel Prize in Chemistry.

These apparent successes notwithstanding, it should be noted that for several reasons the accuracy of our predictions and/or the length of the time that elapses between the signal and the breakthrough could differ for setB relative to the gold standard or setA signals. First, the exponential expansion of the PubMed database since 1997, which has continued through the present day, may explain the narrower range of %New values (compare y-axis values in panels B,D with those in A,C in Fig. 4), and could reduce accuracy by lowering the signal-to-noise ratio (compare G with E in Fig. 4). Second, the shift in setB to more clinical topics, which typically involve studies that include human subjects, may result in slower progress, a lower probability of achieving the predicted breakthroughs, or both. Finally, the hypercompetitive environment faced by biomedical scientists has intensified over the past few decades, raising the risk that a nascent topic might fail to thrive due to insufficient resourcing. Nevertheless, our results overall suggest that information mined from the CCN/RMCL network can be used to predict which scientific topics will produce breakthroughs well in advance of their occurrence.

## Discussion

The potential use of citation networks to track the emergence and association of ideas in science was first recognized in the 1950s and 1960s (*52–54*). In parallel 1973 landmark studies, Irina Marshakova Shaikevich and Henry Small independently described the construction and use of CCNs to cluster publications focused on specific topics (*19, 55*). We began by taking a similar approach but at scale, building a series of 37 CCNs that capture the entire PubMed database through the end of each year from 1981 to 2017, and using the RMCL algorithm to map the topic clusters in each of those networks. Linking each cluster to its counterparts in adjacent years captures the longitudinal progress in that area of research by creating a trajectory for each topic; collectively those trajectories map the progression of biomedical knowledge in the published literature over time. This approach therefore has the potential to realize Gene Garfield’s vision of a dynamic “Atlas of Science” capable of mapping the scientific landscape and detecting the emergence of new scientific fields in real time (*56*).

Scientists have long been drawn to areas of research they judge likely to produce significant discoveries (*57*); our work provides another data point researchers may consider against the background of everything else they know. Some practitioners, if they perceive the opportunity to contribute to a significant discovery, might be persuaded by this additional information to shift their focus, particularly if their current work is in a prediction-adjacent area. Increased awareness could also help organizations optimize the potential for their portfolios to share in major scientific achievements. In this way, both the approach described here, and a related approach to predict FDA approvals described elsewhere (*58*), could both identify promising areas that might otherwise be underappreciated, and accelerate progress in developing new therapeutics.

Balancing these potential positive outcomes is the risk that an over-exuberant pursuit of topics with breakthrough signals will draw resources away from any number of critically important areas of research where the signal of a coming breakthrough has yet to appear. This is a valid concern and highlights the importance of using a multifaceted set of criteria in decision-making, striving to recognize the full range of contributions to scientific progress (*59*). Beyond failing to adhere to this very important precept, other factors seem likely to limit the extent to which the use of our predictive AI/ML approach might lead to maladaptive changes in the scientific ecosystem. Changing research directions is time and resource intensive in all but the earliest stages of a career, and personal connections to a particular problem can further decrease the likelihood a practitioner will change fields (*60, 61*); many funders and publishers likewise focus on a narrowly defined mission, often a particular disease or condition. Though these seem to be the most likely factors involved, it would be unwise to speculate, either how the scientific community will react initially, or how individual scientists might balance the benefits and risks of using this new information over time.

Models of scientific publication, communication, and funding have continuously evolved over the decades, and this process continues in our current era. In particular, open access publishing, data sharing, and the volume of science deposited on preprint servers has grown over the past four or five years. Like the future actions of scientists, funders, and publishers, these changes could theoretically impact our ability to detect future breakthroughs in ways that are difficult to anticipate, and might necessitate periodic updates to our methodological framework. An obvious direction for future work would be the incorporation of preprints into our CCNs, since multiple studies have shown that peer review by journals introduces relatively small changes in reported results (*62–65*). Such an addition might lead to earlier detection of breakthrough signals without any reduction in accuracy, though the technical challenge in reconciling the reference lists of two versions of the same research report (preprint and subsequent peer-reviewed publication) is non-trivial. Finally, since the signals identified by our approach appear to reflect patterns in the behavior of practicing experts, barring unexpectedly rapid changes in those patterns, we anticipate that the basic framework described here will continue to be useful going forward.

The success with which we identified past winners of either a Lasker Award or a Nobel Prize in Medicine that were not included in our training set of gold standards (table S3) suggests that at worst we have detected a majority of the breakthroughs in the time frame of our analysis.

Notably, we also identified behavioral and social science topics in both setA and setB, despite the fact that very important advances in these areas of research are less likely to receive a major prize (*66, 67*). This is not to suggest that our results are free of false negatives, i.e., we are not claiming to have identified all upcoming transformative discoveries in biomedical research. Slow-growing fields with a small number of practitioners may represent special cases that require extra analysis to validate (e.g., aDNA; table S3). For these reasons, and in general as part of best practices, both the positive and negative results of this and similar future work should never be taken at face value, but instead must be “flagged for inspection” and interpreted by subject matter experts in the broader context of a diverse set of parameters.

On the surface, the predictability of many breakthroughs may seem in conflict with the commonly accepted role of serendipity in discovery (*68–71*). However, a successful breakthrough requires a step beyond the unexpected revelation—its recognition by other practitioners, beginning with those closest to the discovery. This recognition can actually precede the moment of serendipity; in such circumstances, scientists start out with shared confidence that the solution of a particular problem will be transformative, and proceed to pursue the breakthrough aggressively (*72*). Whether recognition comes before or after a discovery, the precise nature of the corresponding insight, as well as the circumstances surrounding it, are inherently difficult or impossible to predict. Regardless, we show here that scientists’ collective pursuit of research that will go on to produce breakthroughs generates regular patterns in the CCN/RMCL data that have predictive power. In this way, our work identifies a potential common mechanism for recognizing scientific advances across all domains of biomedical research.

The signals of future breakthroughs have a shared combination of features: a burst of papers exploring a new scientific concept, an unusually high number of very influential papers, and low topical cohesion of the associated content. The most likely explanation for the burst of publication and citation activity is excitement about the potential for a major advance in scientific knowledge. This has been described as the concept of “multiples,” that a breakthrough in a particular topic is “in the air” (*68, 73, 74*). In fact, the combined contribution of numerous researchers may be essential to progress; the initially low topical cohesion can be thought of as an expansion of the opportunity space that occurs when scientists use their different perspectives to explore an exciting new avenue of investigation. Consistent with this framing, a previous analysis of the network of people, institutions, and publications that contributed to the development of two revolutionary new therapies—for cancer (the immunotherapeutic agent ipimilumab, one of the gold standard in this analysis), and for cystic fibrosis (ivacaftor)−concluded that a large, collaborative, and diverse set of investigators is required to support major advances in medicine (*75*).

A parallel, non-mutually exclusive way to explain the formation of the diagnostic signals of future breakthroughs is as productive forays into the adjacent possible (*76–79*), where one or more new ideas on a topic are recognized as most promising because they are adjacent to what is already known in that area of research (commonly used terms like “state of-the-art” and “cutting edge” come to mind). Clusters displaying the diagnostic signal typically appear de novo, as new papers on a shared topic extend the existing scientific predicate of relevant older papers, and are often focused on a dynamic and entirely novel idea. Scientists studying an area that looks particularly promising are likely to explore that adjacent possible/novel idea, and when successful produce papers that are highly influential and trigger the process of transformative emergence. The subsequent breakthroughs, all of which appear to attract outsized levels of interest, display different CCN patterns, either splitting off as a distinct topic (divergence), merging with a previously separate line of investigation (convergence), or maintaining the topic’s central theme (progression).

Each year only a few of the tens of thousands of topics display a diagnostic signal of a future breakthrough; on average those signals appear more than 5 years in advance of the publication(s) announcing the discovery. Whether or not all of the setB topics eventually produce a breakthrough, the work described here suggests a worthwhile framework for making accurate predictions going forward. Future attempts to solve this problem should benefit from further investigation of any false positives; such analytical “autopsies” of those failures could suggest alterations that would improve performance. A large enough collection of successful predictions should eventually allow a more granular level of forecasting, i.e., predictions about the precise nature of the future breakthrough, which in turn might suggest the most promising lines of investigation within each emerging topic.

As with any predictive models, great care must be taken during their construction to ensure that their output is as free of bias as possible; several of our design choices in the work described here are intended to mitigate against this possibility. First, we have incorporated data from all journals indexed in PubMed, since limiting to a subset might unintentionally skew the results toward or away from particular topics. Our data confirm the importance of casting a wide net; specialty journals with a smaller audience, rather than those that are high profile and/or with a broad multi-disciplinary scope, are the most common venue of publication in clusters that signal a coming breakthrough. Second, we have avoided journal- and author-based inputs because they present multiple avenues for the introduction of bias. High visibility journals are not equally accessible to all fields or to all researchers, and cumulative advantage, also known as the Matthew effect (*80*), may be a major driver of results when either journal- or author-based metrics are used. In fact, the agnostic means of identifying promising topics at scale described here might provide a useful check on blind spots and potentially avoid delays in recognition of important scientific advances; recent evidence suggests that major awards map to a relatively narrow range of topics (*81*).

Several lines of further exploration of the work presented here have the potential to improve our understanding of the structure of the scientific enterprise. Further studies may shed light on the factors that differentiate breakthrough and near-breakthrough topics. They might also explain how allocation of resources, workforce dynamics, the stochastic nature of scientific insights, serendipity, and other factors contribute to the advancement of the research enterprise. Finally, since this approach is transposable to citation networks of any size or composition, it should be expandable to permit detection of breakthroughs in any and potentially all areas of science.

## Acknowledgments

We thank Jim Anderson and the NIH Office of Portfolio Analysis team for thoughtful feedback on an earlier version of this manuscript, Grant Jones and Abbey Zuehlke for help with figure preparation, and Summer Allen for assistance with manuscript preparation.

## Author contributions

Conceived and designed the analysis: M.T.D, B.I.H, B.L.B., P.M., T.A.H., K.A.W., and G.M.S. Project management/oversight: R.A.M., K.A.W., and G.M.S (with special thanks to Abbey Zuelke and Grant Jones). Data management and acquisition: M.T.D., B.I.H., B.L.B., P.M., T.A.H., and G.M.S. Analyzed the data: M.T.D., B.I.H., B.L.B., P.M., S.A., T.A.H., R.A.M., K.A.W., and G.M.S. Tool/code development: M.T.D, B.I.H, B.L.B., P.M., T.A.H., and G.M.S. Contributed to the writing/editing to the paper: M.T.D., B.I.H., B.L.B., R.A.M., K.A.W., and G.M.S. This work was performed prior to T.A.H joining the Centers for Disease Control and Prevention and should not be considered a research product of that agency.

## Data and materials availability

All data needed to evaluate the conclusions in the paper are present in the paper and/or the Supplementary Materials. Additional data related to the paper may be requested from the authors.

## Supplementary Materials

Materials and Methods

Figs. S1 to S17

Tables S1 to S3

Movies S1 to S4

Data S1 to S7

Supplementary Text

