## Supplemental Figures, Tables, and Text for "Prediction of transformative breakthroughs in biomedical research"

**This file includes:**

Materials and Methods
Figs. S1 to S17
Tables S1 to S3
Captions for Movies S1 to S4
Captions for Data S1 to S7
Supplementary Text

**Other Supplementary Materials for this manuscript include the following:**

Movies S1 to S4
Data S1 to S7

### Materials and Methods

#### Data sources

Publication and citation data were extracted from the NIH Open Citation Collection (NIH-OCC) for articles published between 1981 and 2017 (as of February 2019). The NIH-OCC comprises the entire PubMed database and includes citations from PubMed and CrossRef, as well as those extracted from the full text of open access articles, using the machine learning algorithm described in Hutchins et al. 2019 (*82*); these data are publicly available through the *iCite* tool (*83*). Review papers were identified as those matching the "Review" flag in the MEDLINE/PubMed Data Element (Field) Publication Type (PT) field (*84*). The complete frozen dataset and associated metadata used to generate the results reported here can be found at the NIH Figshare Archive (https://figshare.com/s/ca05c0539e77fb2a1dc1); sample code for generating RMCL clusters, their associated trajectories, and the identification of breakthrough signals is available in a dedicated Github repository (https://github.com/NIHOPA/predictivebreakthroughs).

#### Generation of co-citation networks and RMCL clusters

The 18.5 million papers that have been deposited in the PubMed database through the end of 2017 are the starting dataset for our analysis. We computed the co-citation network (CCN) of each of those 18.5 million papers, using first-order approximation, for articles that are co-cited with another article in the network. Each paper is represented as a sparse vector that contains a variable number of elements; each element represents a co-cited paper, the value of which equals the number of times it is co-cited with the paper described by the vector. An example is given in Supplementary Text. Briefly, the cosine similarity of all 18.5 million vectors are calculated pairwise against each other, and the resulting values are assembled into a matrix where papers are connected by the cosine similarity weights of their CCNs. All edges with a cosine similarity greater than 0.35 were included; see Supplementary Text for more details. As a paper is increasingly cited, the number of elements in the vector increases, as does its magnitude; using the cosine similarity values rather than the value of the vectors themselves alleviates to some extent distortions that might be introduced when the subject matter of two papers is essentially identical, but one has been more heavily cited than the other.

The above-described matrix can be represented as a network graph, as shown in Figure 1D. We assembled thirty-seven independent graphs, each of which contained no papers, citations, or other data beyond the terminal year x, where x equals a year in the range 1981-2017. Computation of these historical networks allowed us to track citation dynamics through time without leaking information from the future. For the CCN that includes all articles through the end of 2017, this requires 14.7 billion computations at ~500k/second, the equivalent of approximately eight hours run time on an Amazon Web Services (AWS) EC2 server with 96 cores. The smaller number of papers in each of the additional thirty-six earlier historical networks required proportionally less compute. The generation of all thirty-seven networks required approximately 80 hours of processing time in total.

We next applied the Regularized Markov Cluster (RMCL) algorithm (*33-35*) to these thirty-seven CCNs to identify the high betweenness edges that bridge distinct topics. We settled on this method after testing several other methods of building and clustering networks (fig. S1-S4, and Supplementary Text). To prioritize the maintenance of weak connections and reduce the risk of over-splitting topics, we chose a low inflation setting (1.20) for the algorithm (see **Supplemental Methods**). For the most recent and therefore largest CCN (all articles 1981-2017), an inflation setting of 1.2 required approximately four hours of processing time on an AWS EC2 server with 96 cores; RMCL analysis of all thirty-seven networks at an inflation setting of 1.2 required approximately 60 hours of processing time in total.

Using the above parameters to process the co-citation network for all years included in our dataset (1981-2017), RMCL designated 339M intra-cluster edges and 39M inter-cluster edges (a median of 17 and 2 per publication, respectively) within the CCN, thereby distributing 17.2M publications across 23,867 discrete topic clusters. The remaining 1.3M publications (7.1% of the total), which had few citations (a median of 1) and few CCN links to other papers (a median of 1 intra- and 1 inter-cluster edge per publication), were not considered further.

Networks were visualized by collapsing the collection of publications in each RMCL cluster into one node. To allow visualization in Cytoscape (*85*), the edges between clusters were collapsed into a single edge if the summed edge weight was 25 or higher. Edges that did not sum to the threshold for summed edge weight were not included.

**Content analysis of RMCL clusters**

We quantified the semantic similarity of articles assigned to distinct RMCL clusters using a well-established language model, word2vec (*36, 37*). The content of each article was represented by its title and abstract, the text of which was concatenated and preprocessed using our text pre-processing library NLPre, which corrects for inconsistencies in real-world data such as random capitalization patterns, strange hyphenations, and abbreviations (*36, 86*). To more efficiently manage the large size of the embedding, processing speed was improved by using gensim (*87, 88*). Document vectors were constructed by summing each word vector weighted by its inverse document frequency. Cluster level centroids were constructed by summing all document vectors and a representative set of words was chosen by taking the closest words in the embedding space to the centroid vector.

To evaluate the alignment between RMCL assignment to clusters and established methods for defining topics, research articles published between 1995 (the first year for which data on article type is available) and 2017 were drawn from six leading journals: three multi-disciplinary (*Nature*, *Science*, *PNAS*) and three disciplinary (*Blood*, *Genetics*, *Journal of Neuroscience*). Article types other than research articles (e.g., reviews, commentaries, and other front matter) were excluded. A total of 9,147 articles published in Genetics met these criteria, all of which were selected for analysis; 10,000 articles were selected at random from each of the other five journals. 100k pairs of articles were randomly drawn from each journal-specific dataset; the word2vec cosine similarity of each pair's semantic content was computed, and their similarity distribution was analyzed on a per-journal basis. Across all journals, 2364 article pairs shared both their venue of publication and their RMCL designation. These pairs' similarities were also analyzed as a separate subset. As a control we also drew pairs of articles that were not published in one of our six journals, but had been assigned to one of the clusters flagged by those journals. 100 RMCL clusters containing at least 1000 publications (to reduce the odds of duplicate draws) were chosen at random from the set of clusters containing at least one pair of articles from the previously described journal-specific data. 10,000 pairs of articles were then drawn from each of these clusters, excluding anything published in one of our six journals, resulting in a combined 100k pairs whose semantic similarities were likewise computed. As an additional control, the process was repeated using the set of n clusters that contained no publications in our journal dataset.

**RMCL versus coding by subject matter experts**

We determined the level of agreement between RMCL clustering and human judgment by asking subject matter experts at NIH to partition publications assigned to pairs of RMCL clusters with increasing word2vec similarity (*36*). Five experts were provided with 10 publications from two RMCL clusters (five from “group A” and five from “group B”) and were permitted to use titles and abstracts to sort the articles into two groups. The partitioning experiment was repeated for ten pairs of RMCL clusters, whose word2vec similarity ranged from 0.0 (low similarity) to 0.9 (high similarity), so that each human expert evaluated, in total, 100 publications across that entire range; results are summarized in Data S1; similarities for a random sample of 26,330 pairs of PubMed IDentifiers (PMIDs; the unique identifiers assigned to each paper in the PubMed database by the National Library of Medicine) are shown in fig. S14. Average inter-rater reliability between expert judgment and RMCL was calculated to be 0.96 via pairwise Cohen's Kappa testing, indicating that the RMCL clustering aligns well with human judgment.

**Trajectory generation and visualization**

We generated trajectories to track the flow of information from year to year by identifying a cluster's historical counterparts in the year following (x+1; forward direction) and preceding (x-1; backward direction) a given reference year (x). Clusters were designated as historical counterparts if they shared over 50% of their publications; across all years, a historical counterpart could be identified for 98.7% of all clusters. A variety of other, more complex algorithms for linking clusters over time might be devised. However, since perfect 50:50 splits occur less than 0.08% of the time, this simple design choice has the advantage of clearly identifying the single most highly related clusters year over year; any other method would, by definition, link clusters that share only a minority of their papers. To account for rare cases where a cluster's papers split nearly evenly into two descendants (splits in the 45-55% range account for 2.5% of all cases), and to avoid losing information from clusters that might be substantially related despite sharing only a minority of papers, for each reference cluster (x), we tracked the two clusters with the highest number of overlapping PMIDs, in the following (x+1) and preceding (x-1) years using the binary measures described below:

| **Descriptor** | **Temporal direction** | **Definition** |
|---|---|---|
| has_host | Forward | Is there a cluster in year x+1 (i.e., the next year) that contains over 50% of these PMIDs? |
| comprises_host | Forward | Do PMIDs from the year x (i.e., the reference year's) cluster comprise over 50% of the candidate historical counterpart? This identifies the most highly related cluster in year x+1. |
| comprises_host_alt | Forward | Of the clusters remaining after designating comprises_host, do PMIDs from the year x cluster comprise over 50% of the candidate historical counterpart? This identifies the second-most highly related cluster in year x+1. |
| has_ancestor | Backward | Is there a cluster from year x-1 (i.e., the previous year) that contains over 50% of these PMIDs? |
| comprises_ancestor | Backward | Do PMIDs from the year x cluster comprise over 50% of the candidate historical counterpart? This identifies the most highly related cluster in year x-1. |

| | | |
|---|---|---|
| comprises_ancestor_alt | Backward | Of the clusters remaining after designating comprises_ancestor, do PMIDs from the year x cluster comprise over 50% of the candidate historical counterpart? This identifies the second-most highly related cluster in year x-1. |

The two most highly related descendants of a cluster (i.e., comprises_host and comprises_host_alt) include upwards of 90% (mean 90.9%, median 93.4%) of its PMIDs. To capture the remaining <10%, we developed a vocabulary to describe all the different types of links we observed between clusters from one year to the next (listed in the table below; for corresponding schematics see fig. S7). Some were categorized as “major”, indicating links between clusters designated as historical counterparts , while others were categorized as “minor”, tracking related clusters for which there were lower levels of overlap.

| **Link description** | **Temporal direction** | **Link type** | **Criteria** |
|---|---|---|---|
| 1-to-1 | Forward | Major link | has_host AND comprises_host |
| Captured | Forward | Minor link | has_host AND (NOT comprises_host) |
| Split | Forward | Major link | (NOT has_host) AND comprises_host AND comprises_host_alt |
| Shed | Forward | Major link | (NOT has_host) AND (comprises_host OR comprises_host_alt) |
| Disintegrate | Forward | No link | NOT (has_host OR comprises_host OR comprises_host_alt) |
| Bud | Forward | Minor link | has_host AND comprises_host AND comprises_host_alt [this link is made for the next year's cluster with the second largest number of overlapping PMIDs] |
| 1-to-1 | Backward | Major link | has_ancestor AND comprises_ancestor |
| Split (recombination ) | Backward | Minor link | has_ancestor AND (NOT comprises_ancestor) AND comprises_ancestor_alt |
| Split (clean) | Backward | Minor link | has_ancestor AND (NOT comprises_ancestor) AND (NOT comprises_ancestor_alt) |
| Merger (two) | Backward | Major link | (NOT has_ancestor) AND comprises_ancestor AND comprises_ancestor_alt |
| Merger (lots) | Backward | Major link | (NOT has_ancestor) AND comprises_ancestor AND (NOT comprises_ancestor_alt) |
| Minor merger | Backward | Minor link | (NOT has_ancestor) AND (NOT comprises_ancestor) AND comprises_ancestor_alt |
| De novo formation | Backward | No link | NOT (has_ancestor OR comprises_ancestor OR comprises_ancestor_alt) |

A cluster may be in more than one trajectory under specific limited circumstances. A cluster may split from one year to the next so that a single cluster A in year x is the ancestor for two smaller clusters B and C in year (x+1), both of which share a majority (≥50%) of their papers with A. In the absence of a later reunification event, mapping the trajectory of B and its progeny or C and its progeny backwards in time results in two independent trajectories. Alternatively, multiple smaller clusters that are formed by a combination of new papers entering the network and existing papers leaving their previously assigned cluster(s) may share a majority of their papers with one larger cluster.

Trajectories were visualized over the 1981-2017 timeframe as networks in Cytoscape (*85*), using yFiles Hierarchical Layout; human subject matter experts evaluated these visualizations to assess whether each individual trajectory could be classified as convergence, divergence, or progression. Within a trajectory, size is shown as relative to the cluster with the highest number of publications and heatmapping indicates the percentage of new publications relative to the cluster with the highest value. Minor links are included in visualizations for completeness, but were pruned prior to further quantitative analysis.

***Statistical analysis of the gold standard trajectories***

We observed that the frequency of convergent and divergent trajectories in the set of gold standards is high relative to expectations. To compute the statistical significance of this phenomenon, we compared the observed percentage of convergence and divergence events to the baseline probability of those events (0.90% for convergence and 1.47% for divergence); both were statistically significant ($P<0.001$; binomial test).

**Logistic regression**

***Model selection***

We used a logistic regression approach trained on historical data to detect transformative breakthroughs. Logistic regression is a tool that has been applied to similar binary classification problems since the nineteenth century (*89*). It functions by fitting a sigmoid, or "logit" curve, to each independent variable—in our case a set of cluster features (see below)—and combining them to model the probability the test cases will fit into either category. We used Scikit-learn's logistic regression implementation (sklearn.linear_model.LogisticRegression); the default l2 regularization and "lbfgs" solver were sufficient for our purposes (*90, 91*).

For each of the 21 gold standards we used SMEs to identify the cluster most likely to foreshadow the later breakthrough. Notably, the feature those selections had in common was a high percentage of recent papers (table S1). As negative controls we identified the ten clusters closest in size (total number of papers) to each gold standard cluster, five larger and five smaller, all falling within +/-0.03% of the size of the corresponding gold standard cluster.

***Feature selection and vectorization***

We began by generating a list of features that could be calculated for any given cluster in a given year, that is, without relying on information that would have been unavailable beyond that given year. We then narrowed our list to only those features that were statistically distinguishable between gold standards and negative controls ($p< 0.05$ using an independent t-test; see Data S7 for the full list of features we considered). Finally, we conducted an iterative feature elimination experiment, by removing one feature at a time from regression models trained on different combinations of those features listed as statistically distinct in Data S7. In all cases, one hundred rounds of training were performed against a withheld fraction of the training set. From this we identified four features that offered the optimal combination of predictive and explanatory power:

**%New:** The fraction of papers in a cluster that were published less than a year before the date of the cluster (i.e., the percentage of all papers in a cluster that appeared no earlier than year n-1, where n is the year of the cluster).

**ancestral_%New:** The %New of the cluster's largest ancestor (i.e., the percentage of all papers in year n-1 that appeared no earlier than year n-2). Combined with %New, this allows for direct measurement of a cluster's year-over-year growth.

**ancestral entropy:** The fraction of a cluster's papers which were not new, and did not originate in the cluster's two largest ancestors. This measure is highest in clusters that integrate material from multiple fields (i.e., they may be described as interdisciplinary).

**hRCR top10:** hRCR value of the paper at the 90th percentile of all values within the cluster; a measure of how impactful the cluster's papers are beyond the single most highly-cited paper. To avoid leaking information from the future, we computed historical RCR (hRCR) by considering only publications and their citations in or prior to the year of the cluster (i.e., for each paper in year n includes in the calculation only publications and citations that appeared in year n or earlier).

We theorized that a trajectory, which represents the historical track record of a topic, might contain important predictive information about its future development. However, it would be impractical to provide the logistic regression with each of the above four features for all of the ~790,000 clusters that are represented in our dataset of 37 historical networks ending between 1981 and 2018. To retain some hysteretic information about the breakthrough cluster and its trajectory while maintaining a tractable number of features, we therefore calculated the mean and standard deviation value of %New, ancestral_%New, ancestral entropy, and hRCR top10 for the clusters in the five years (years x-1 through x-5) preceding the year of analysis. We then combined these eight values, which represent the central tendency (mean) and annualized variance (standard deviation), respectively, of the recent history of a topic, with the four values specific to the most recent cluster year (x) to generate a 12-number array that served as the input for logistic regression. For example, for the Apoptosis/Bcl2/Bax gold standard, we started with the values for %New, ancestral_%New, ancestral entropy, and hRCR top10 for RMCL cluster #4032 in 1987 and combined them with the mean and standard deviation of the values for %New, ancestral_%New, ancestral entropy, and hRCR top10 for clusters in its trajectory from 1982 through 1986.

To test the robustness of this diagnostic signal, we applied the logistic regression to all clusters in two distinct 4-year windows that are 20 years apart (setA: 1994 through 1997 and setB: 2014 through 2017). This equates to a total of 45,466 clusters for the 1994-1997 window and 140,983 clusters for the 2014-2017 window. Each of these 186,449 clusters was assigned a value of either 1 or 0, indicating they either matched (1) or did not match (0) the profile of a breakthrough, based on the four features extracted from the gold standards. 154 clusters (0.34%) in the set A 1994-1997 window and 705 clusters (0.5%) in the set B 2014-2017 window were assigned a value of 1. All 859 candidates from the two four-year windows were filtered further by selecting the subset of clusters that both experienced recent growth, defined as at least 15% of papers being new in the last year, and displayed a broad distribution of scientific influence. The latter is defined as all papers in the cluster having a historical RCR (hRCR) that was at least average (equal to 1.3), and at least 1/3 of papers having an hRCR equal to or better than the overall median (1.0). Finally, to avoid trivial or non-biomedical positives in the outputs, we limited the analysis to clusters with 40 or more publications, at least 5% of which were linked to an NIH award. After these five filters were applied, 41 and 163 candidates remained in the set A 1994-1998 and set B 2014-2017 time frames, respectively.

Some of these trajectories contain a signal in multiple years; in those cases, we retained only the chronologically first cluster and eliminated later ones. For the 1994-1997 time frame, this left a final 36 candidates. Two of those 36 correspond to Nobel prizes that were announced after our gold standards were chosen (immune checkpoint blockade, 2018 Nobel prize in medicine; and protein engineering, 2018 Nobel prize in chemistry). A further three correspond to our chosen gold standards: SRFM/GFP, 2008 Nobel prize in chemistry; discovery of the genes causing hereditary breast cancer, 2014 Lasker prize; and discovery of leptin-orexin, 2010 Lasker prize. Finally, we eliminated candidates that had either a historical weighted RCR (hwRCR) or a proportion of within-cluster (as opposed to out-of-cluster) citations below the values observed for these confirmed breakthroughs; these two variables capture the extent to which the work was immediately noticed (rapidly became highly cited) and the extent to which it stimulated conversation within, rather than outside, the field. The 16 clusters that remained after these final filters were applied were chosen as the setA breakthroughs. Application of the identical parameters to the 2014-2017 time frame identified 18 predicted breakthroughs (setB).

### Supplementary Figures and Tables

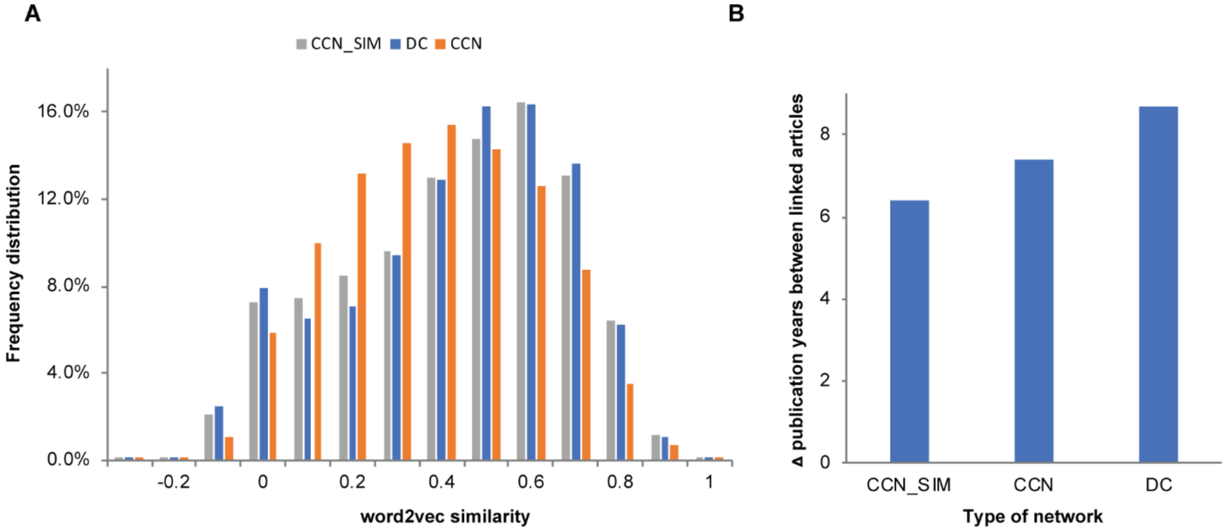


**Fig. S1. Topic and temporal cohesiveness for different network types. (A)** Frequency distribution of semantic similarity as captured by word2vec (*36, 37*) for three types of citation networks: direct citation (DC; blue bars), co-citation (CCN; orange bars), and co-citation with cosine similarity (CCN_SIM; grey bars). The DC appears to outperform the CCN in representing semantic content, a discrepancy that was resolved by using cosine similarities (see Main Text). **(B)** Difference in publication years between linked articles for different types of citation networks: linked articles in CCN_SIM networks are more temporally cohesive than those in CCN or DC networks.

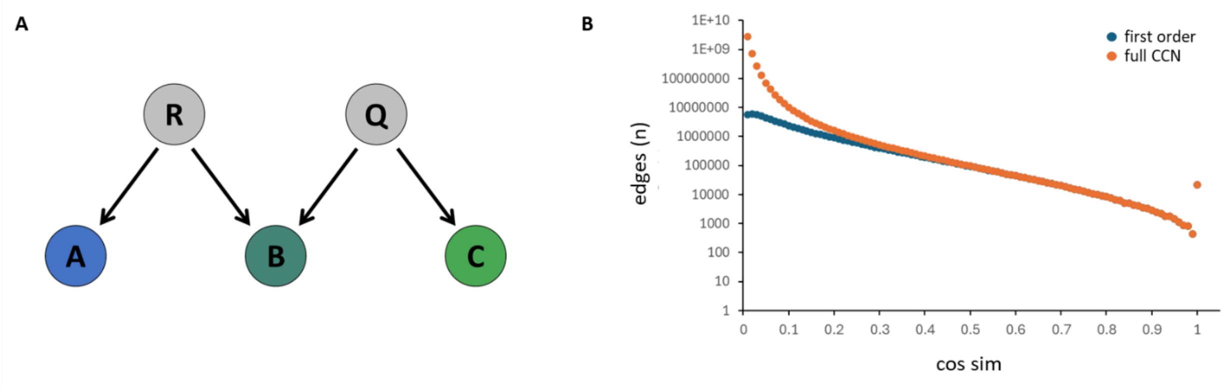


**Fig. S2. First order edges in the PubMed co-citation network. (A)** Definition of first-order edges in a CCN. Gray nodes R & Q represent citing papers; blue and green nodes A, B, & C represent the full CCN generated by R & Q. First-order edges are shown as black arrows. No edge is shown between A and C because they are connected only transitively. **(B)** Cosine similarity (cos sim) was calculated for edges connecting ~100k cited PMIDs selected at random from the full CCN of PubMed. Edges where cos sim < 0.01 are not included. Above a cos sim of 0.35, 93% of all edges in the full CCN are first order edges.

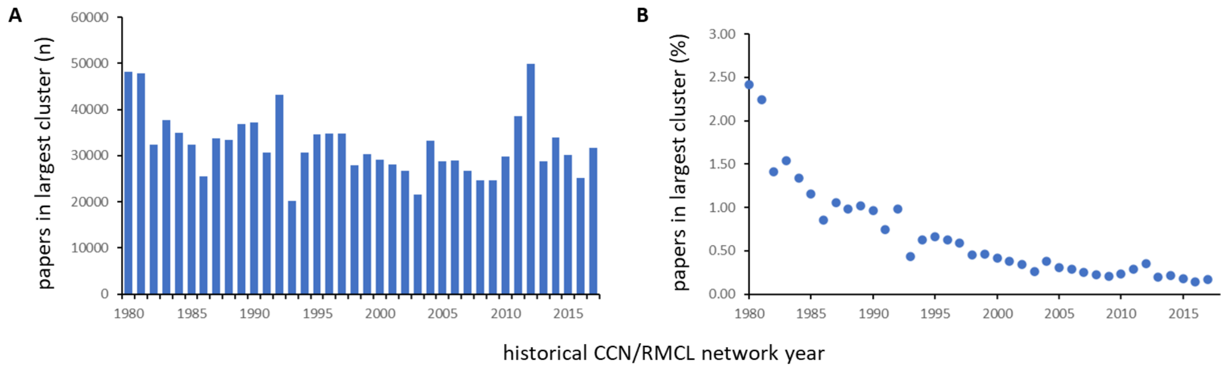


**Fig. S3. Few papers are overmerged in a single large cluster at an RMCL inflation setting of 1.2.** For each historical network, the number **(A)** and the corresponding percentage **(B)** of all cited papers assigned to the largest cluster at an inflation setting of 1.2.

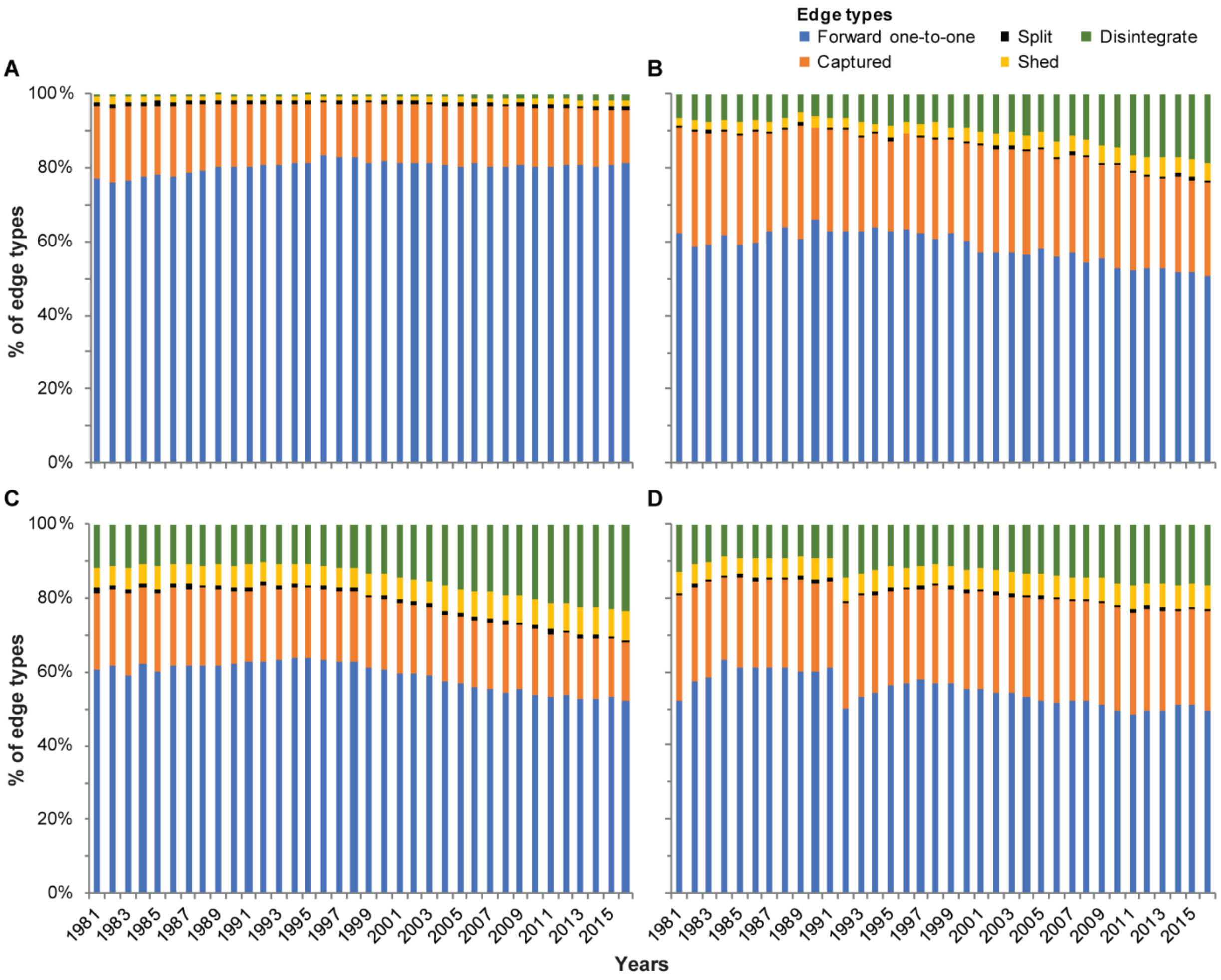


**Fig. S4. Stability of network edges for RMCL compared to other methods**. Stability of network edges (links) were measured over time by tracking the percentage of forward one-to-one links (blue bars), as a proxy for network stability, compared to captured (orange bars), split (black bars), shed (yellow bars), or disintegrated (green bars) links, as a proxy for network dynamicity. **(A)** RMCL applied to CCN edges with an inflation setting of 1.20; **(B)** Leiden algorithm (*93*) applied to CCN edges with a resolution parameter of 0.00020; **(C)** Leiden algorithm applied to CCN edges with a resolution parameter of 0.00001; **(D)** Leiden algorithm applied to direct citation network edges with a resolution parameter of 0.00009. Network stability was maintained most consistently using RMCL at inflation setting 1.20 (panel **A**). See fig. S8, Materials and Methods, and Supplementary Text for schematics and descriptions of edge types.

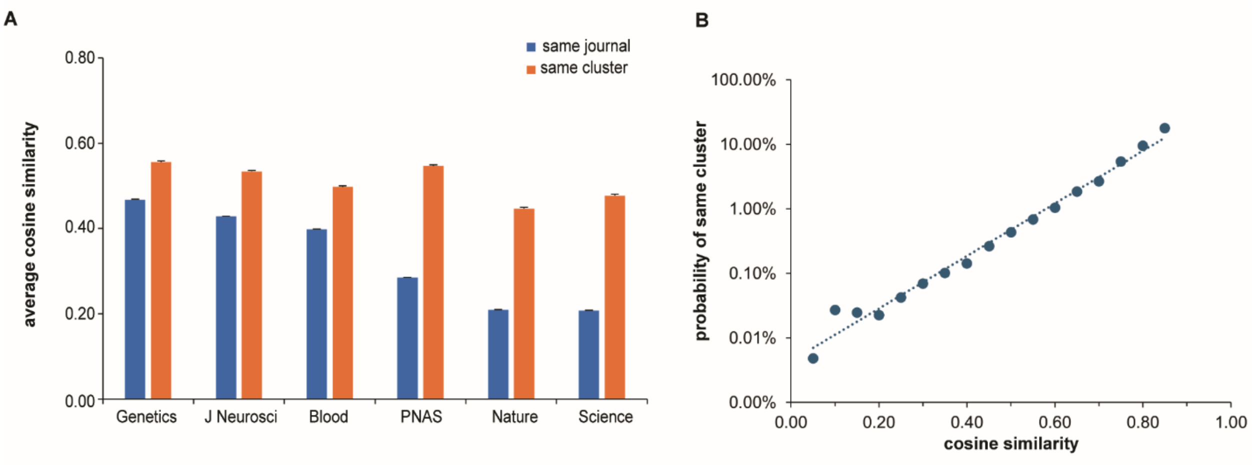


**Fig. S5. RMCL clusters are better at delineating scientific topics than journal of publication. (A)** 50K PubMed articles from the collection of journals shown were paired randomly; pairs that shared the same journal (blue bars) had lower average cosine similarity than pairs that shared the same CCN/RMCL cluster (orange bars). Error bars, SEM. **(B)** As the cosine similarity of randomly paired articles from **(A)** increases, the probability that RMCL assigned both articles to the same cluster increases exponentially.

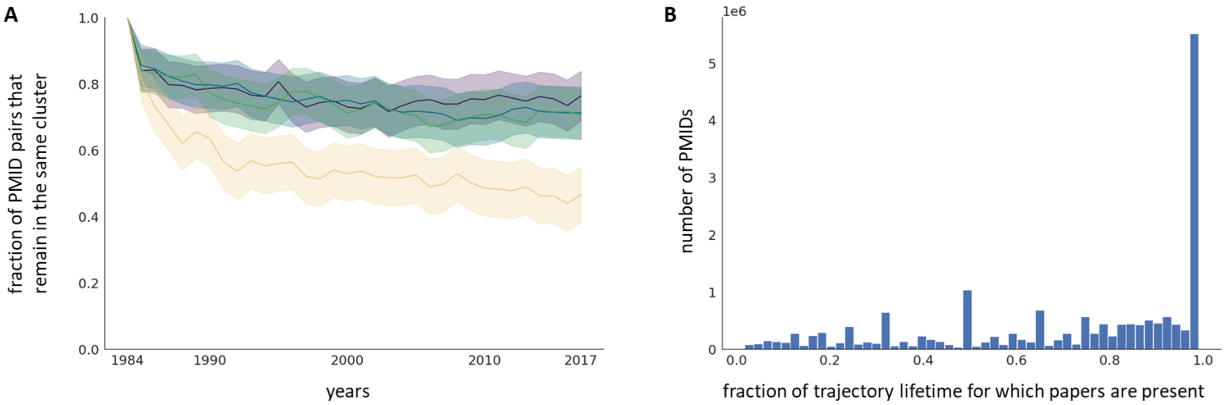


**Fig. S6. Stability of clusters and trajectories. (A)** Assignment of PMID pairs to a shared cluster over time. All papers published in 1981 that received at least 5 citations in the year of their publication were taken as the starting dataset; the 1984 cluster assignment of these papers was identified, and the most semantically similar PMID in that cluster was identified as a partner. The resulting PMID pairs were assigned to quartiles of semantic similarity and followed over time. Purple, top quartile; blue, second quartile; green, third quartile; yellow, bottom quartile. 95% CI, SEM. **(B)** Assignment of PMIDs to the same trajectory over time. Fraction of each individual publication’s “lifetime” spent in the trajectory terminating in its assigned cluster in 2017 is shown; n ≈ 1.8 million PMIDs.

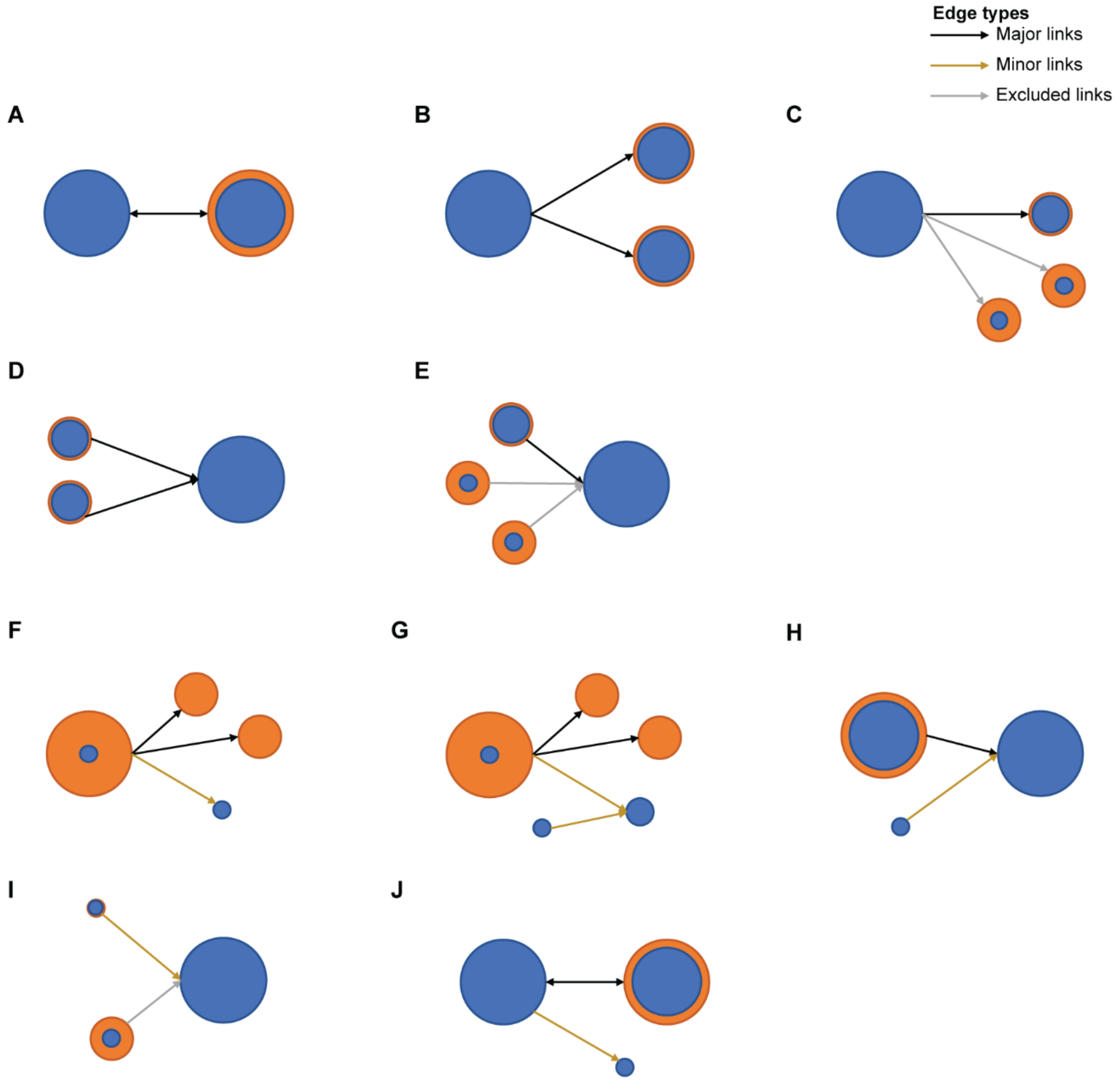


**Fig. S7. Schematic representation of links in trajectories**. Trajectories were generated by connecting clusters in a given year with counterparts in the neighboring years. Blue node shading represents the preserved subset of articles in each cluster from one time period to the next, and orange is the non-preserved subset. Major links between the preserved subset are illustrated in **A-E** and minor links in **F-J** Link types: **(A)** One-to-one; **(B)** Split; **(C)** Shed; **(D)** Merger (two); **(E)** Merger (lots); **(F)** Split (clean); **(G)** Split (recombination); **(H)** Captured; **(I)** Minor merger**; (J)** Bud. See Materials and Methods for detailed definitions of links.

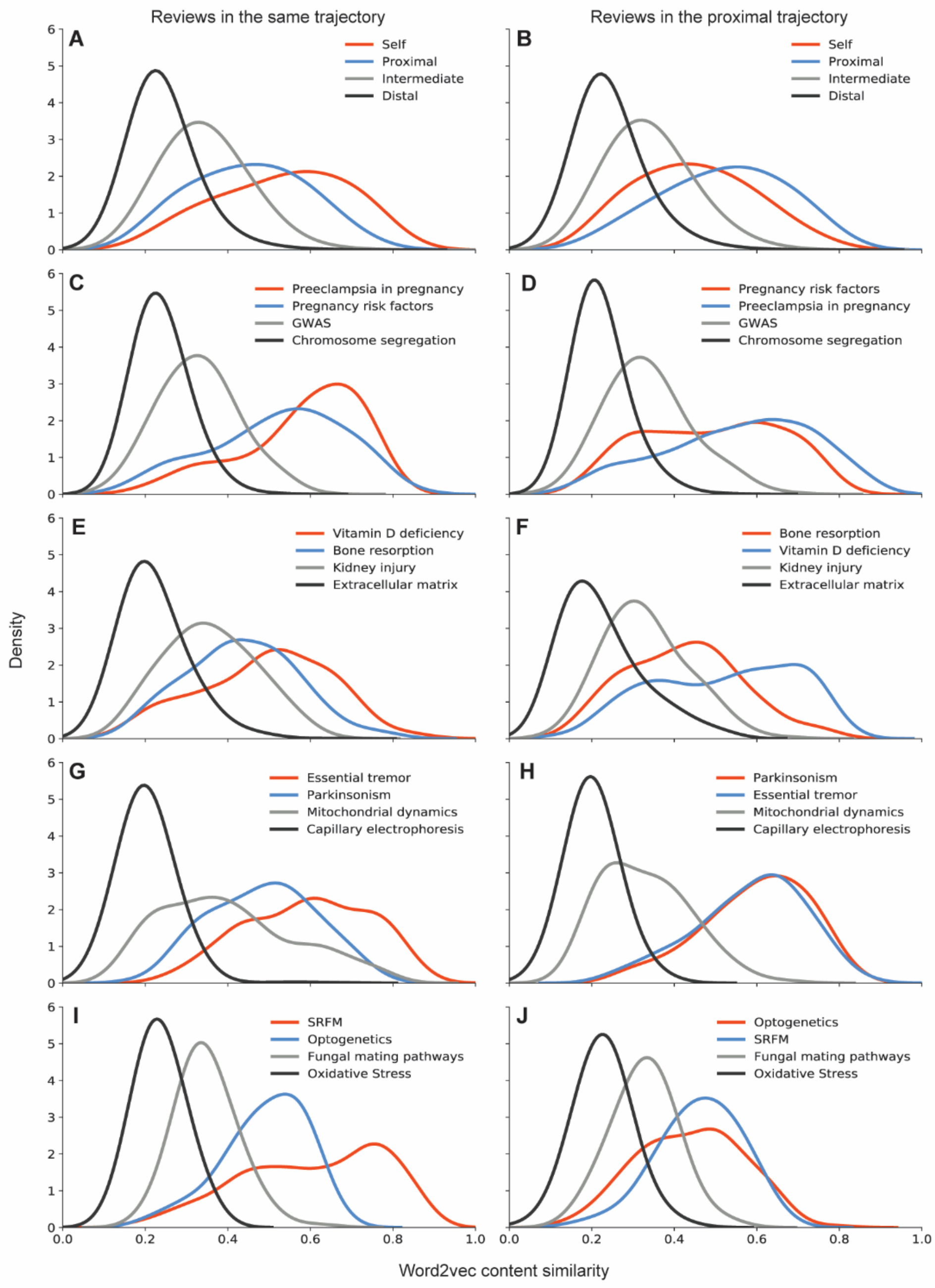


**Fig. S8. Content similarity between reviews and research articles in CCN/RMCL trajectories.** The content in review papers was compared with research reports in the test trajectory (Self; red) or with increasingly distant trajectories in multidimensional space, as measured by word2vec similarity: Proximal (blue), Intermediate (gray), and Distal (black). **(A,B)** 100 randomly selected trajectories in aggregate; **(C,H)** Three of the randomly selected trajectories individually; **(I,J)** the SRFM trajectory; **(A,C,E,G,I)** Similarity to reviews from the Self trajectory; **(B,D,F,H,J)** A reciprocal test, similarity to reviews from the Proximal trajectory.

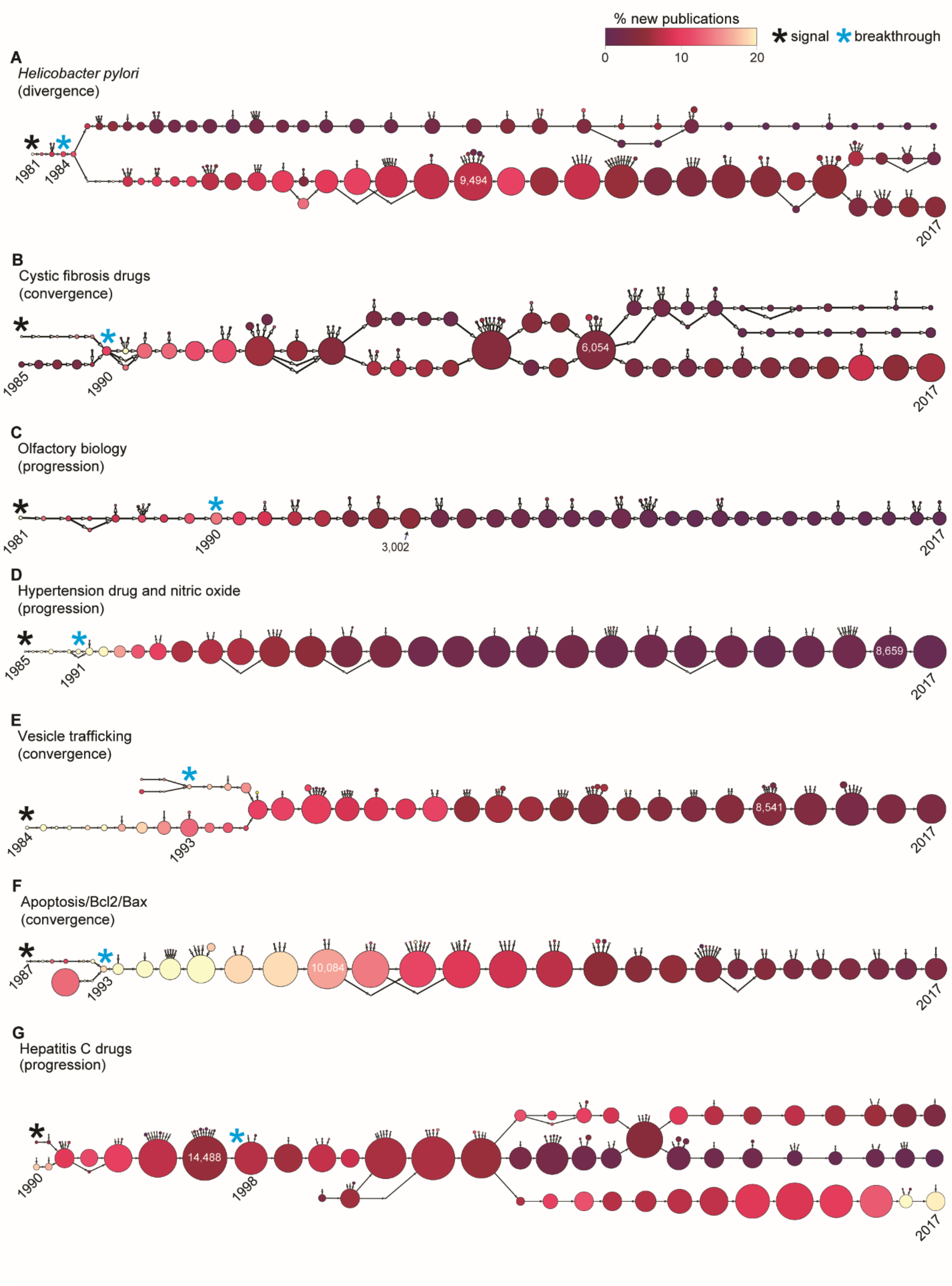
% new publications
0
10
20
signal
breakthrough
A
Helicobacter pylori
(divergence)
1981
1984
9,494
2017
B
Cystic fibrosis drugs
(convergence)
1985
1990
6,054
2017
C
Olfactory biology
(progression)
1981
1990
3,002
2017
D
Hypertension drug and nitric oxide
(progression)
1985
1991
8,659
2017
E
Vesicle trafficking
(convergence)
1984
1993
8,541
2017
F
Apoptosis/Bcl2/Bax
(convergence)
1987
1993
10,084
2017
G
Hepatitis C drugs
(progression)
1990
14,488
1998
2017

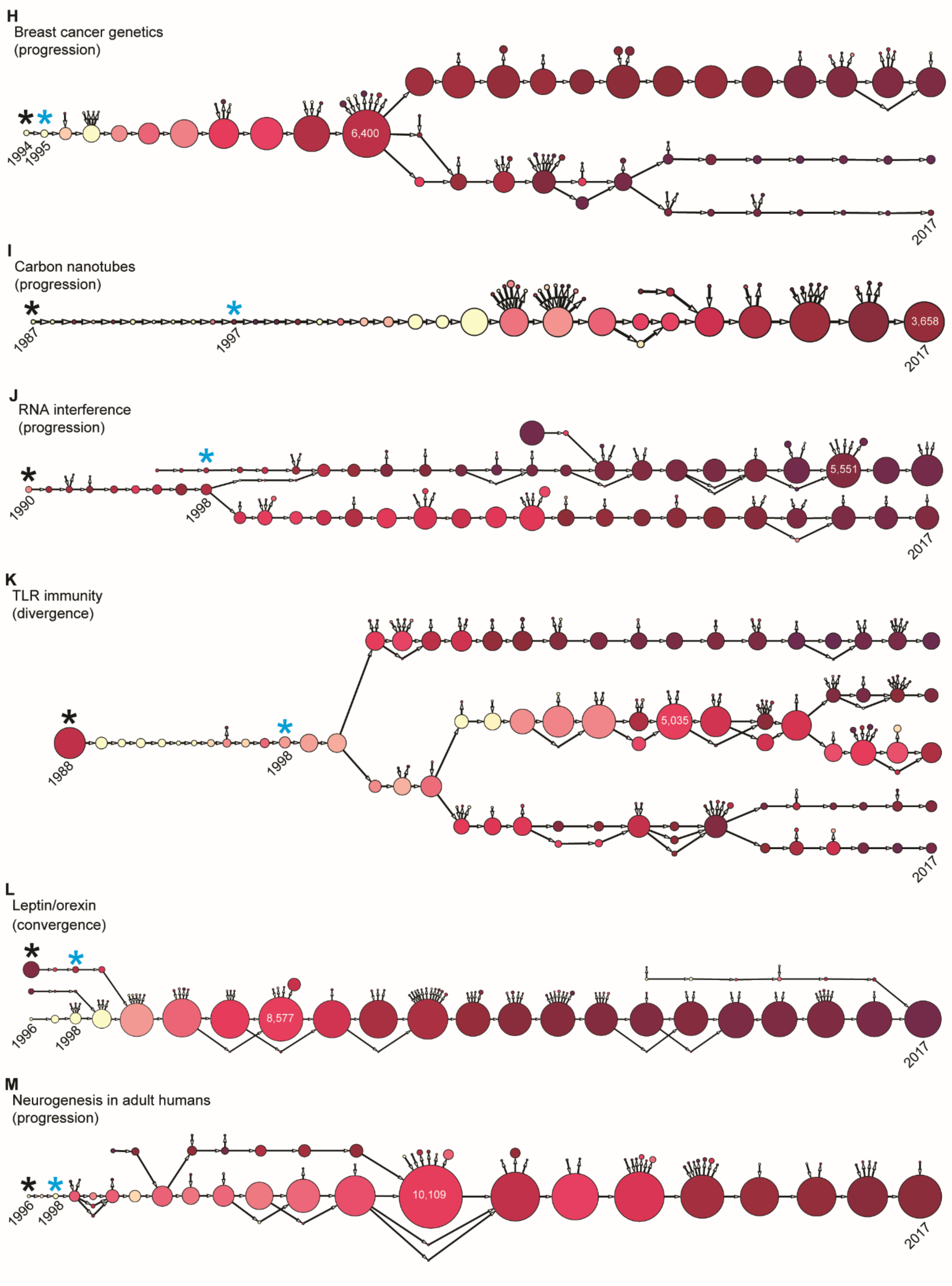
H
Breast cancer genetics
(progression)
1994
1995
6,400
2017
I
Carbon nanotubes
(progression)
1987
1997
3,658
2017
J
RNA interference
(progression)
1990
1998
5,551
2017
K
TLR immunity
(divergence)
1988
1998
5,035
2017
L
Leptin/orexin
(convergence)
1996
1998
8,577
2017
M
Neurogenesis in adult humans
(progression)
1996
1998
10,109
2017

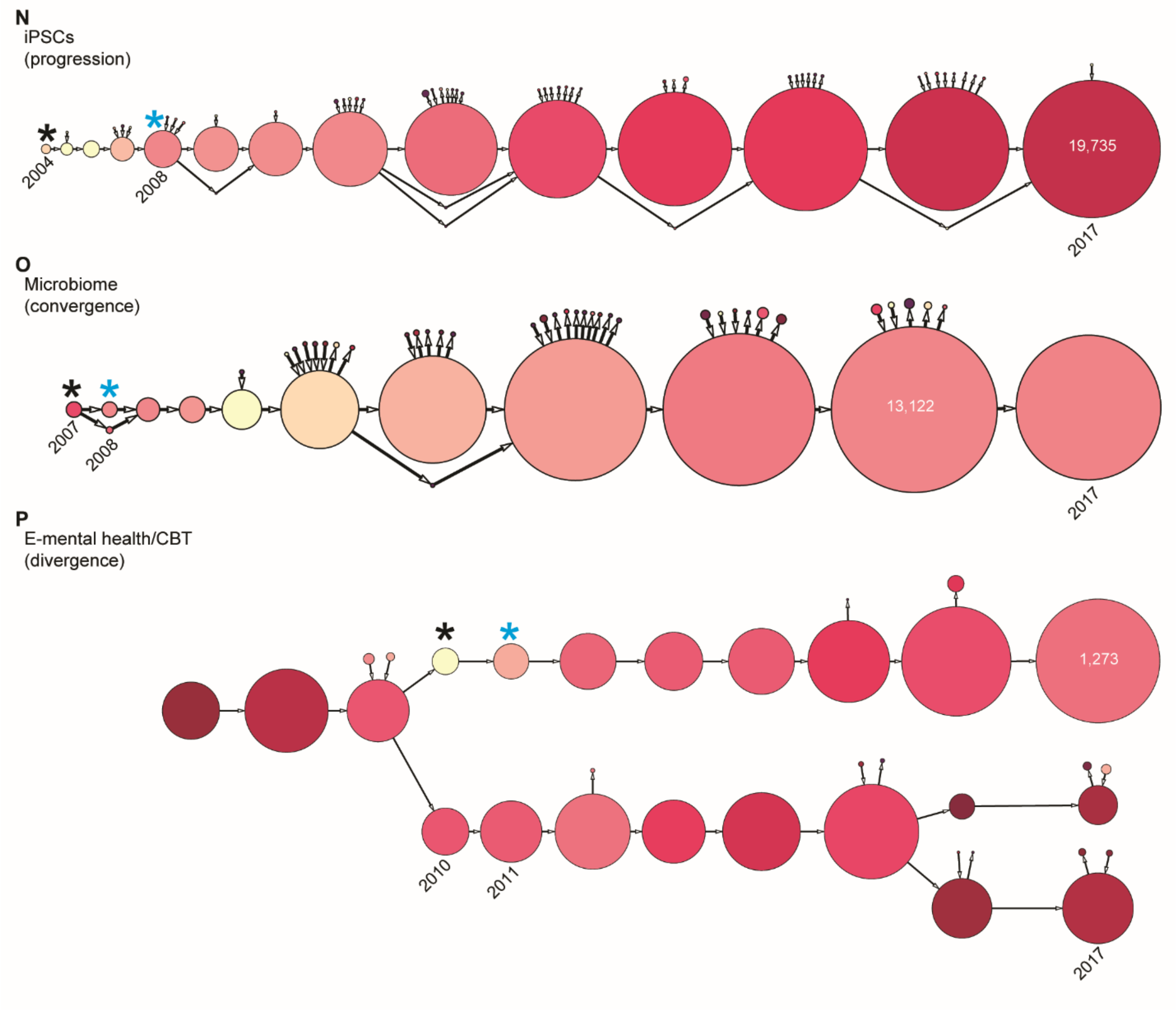


**Fig. S9. Trajectories for gold standard breakthroughs.** Trajectories for the gold standards described in table S1, excluding those that were presented in Figs. 1 and 3. **(A-P)** Nodes are shaded by % new publications and node sizes are relative within each trajectory; the number of articles in the largest cluster for each trajectory is indicated. Signal clusters are denoted with black asterisks and breakthrough clusters with light blue asterisks. Signal year, breakthrough year, and terminal year (2017) is indicated for each trajectory.

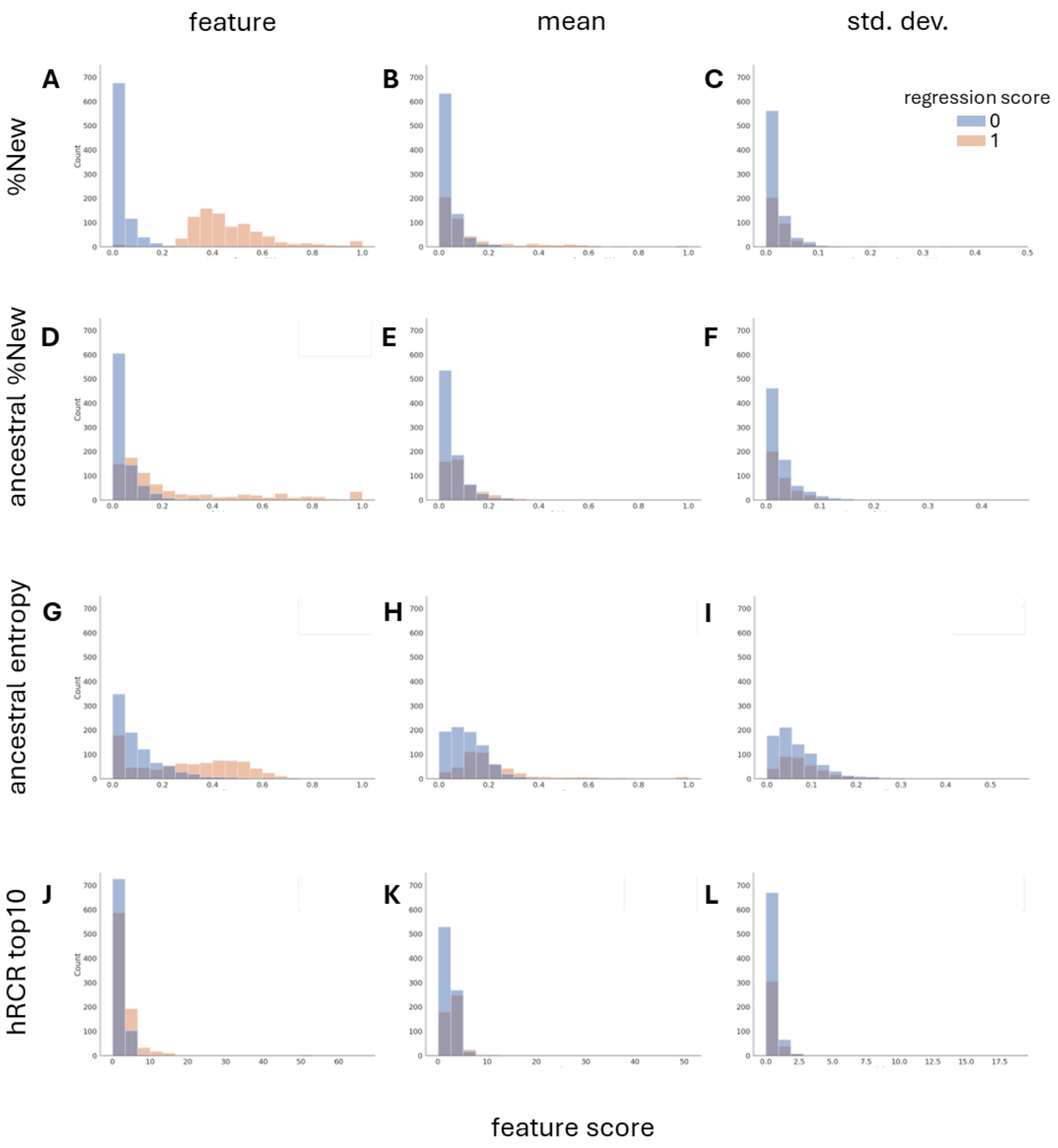


**Fig. S10. Distribution of values for features used in logistic regression.** Histograms of the twelve features computed for each topic and used for logistic regression, separated into negative (0) and positive (1) regression scores. Rows: the four different types of feature computed (see Materials and Methods for details). Columns: Left **(A,D,G,J)** the feature computed for the topic cluster itself in the current year. Center, Right: the mean **(B,E,H,K)** and standard deviation **(C,F,I,L)** of the feature computed for the clusters in the last five years of the topic's trajectory, as described in Materials and Methods.

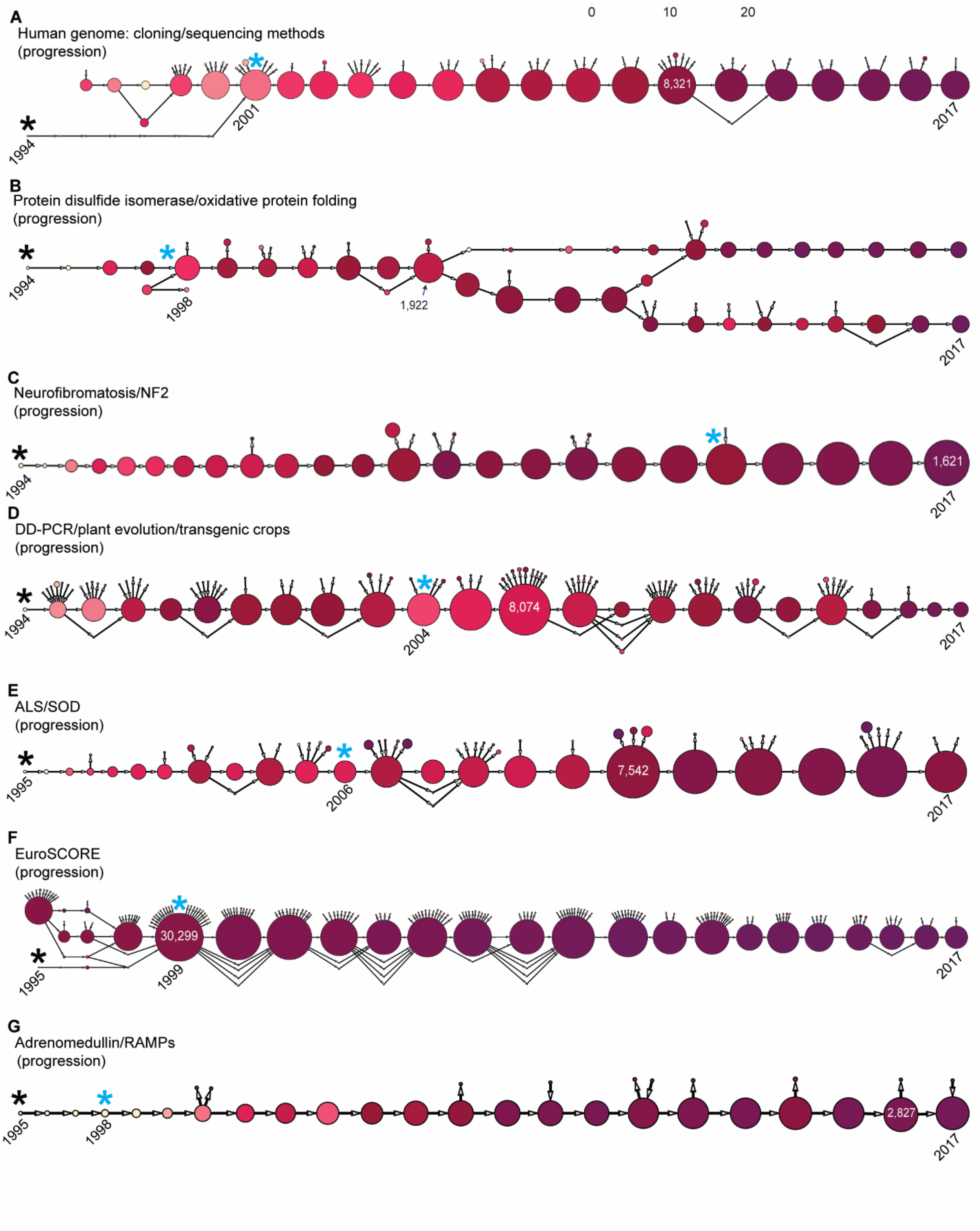
A
Human genome: cloning/sequencing methods
(progression)
0
10
20
1994
2001
8,321
2017
B
Protein disulfide isomerase/oxidative protein folding
(progression)
1994
1998
1,922
2017
C
Neurofibromatosis/NF2
(progression)
1994
1,621
2017
D
DD-PCR/plant evolution/transgenic crops
(progression)
1994
2004
8,074
2017
E
ALS/SOD
(progression)
1995
2006
7,542
2017
F
EuroSCORE
(progression)
1995
1999
30,299
2017
G
Adrenomedullin/RAMPs
(progression)
1995
1998
2,827
2017

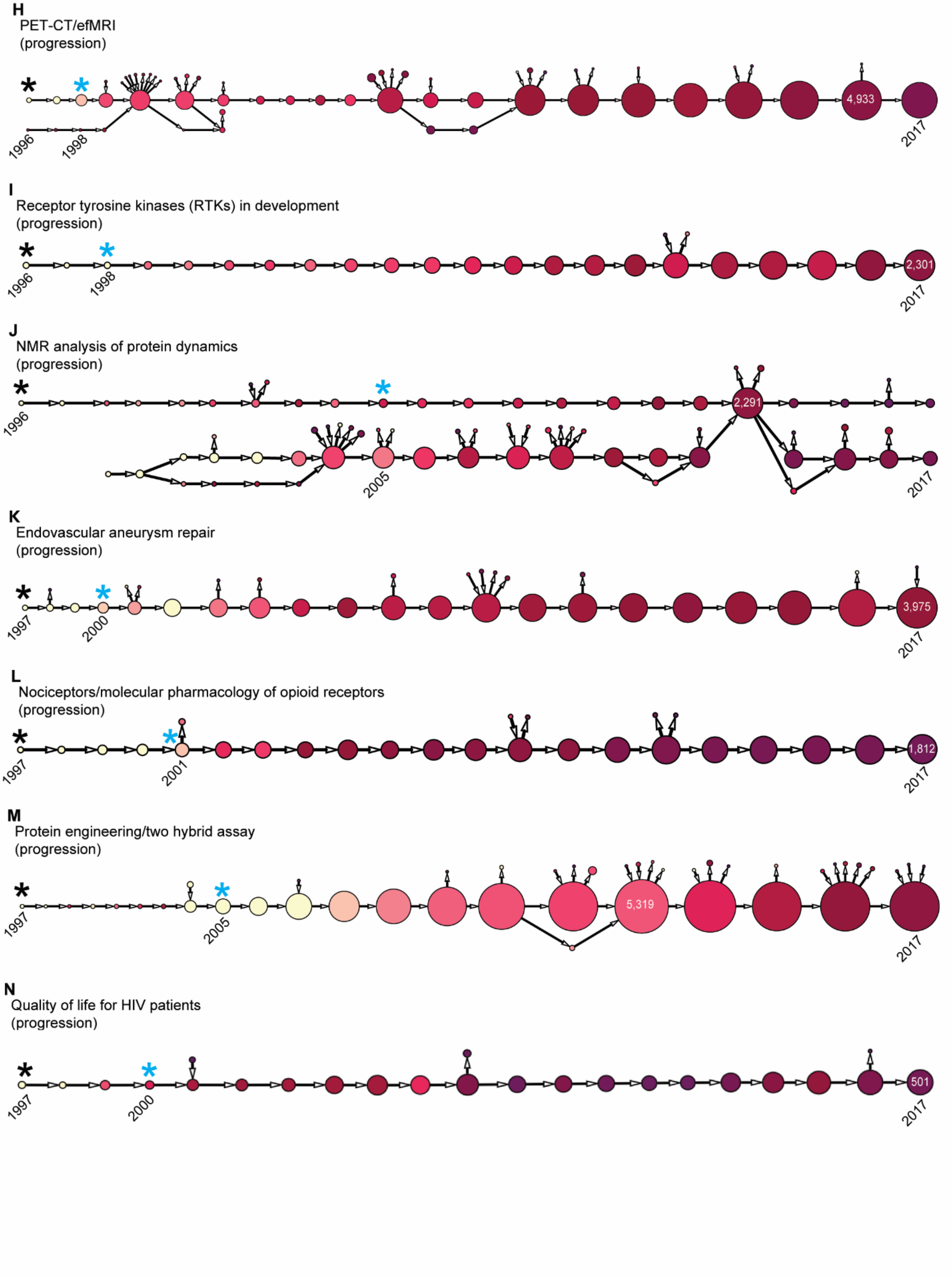
H
PET-CT/efMRI
(progression)
1996
1998
4,933
2017
I
Receptor tyrosine kinases (RTKs) in development
(progression)
1996
1998
2,301
2017
J
NMR analysis of protein dynamics
(progression)
1996
2,291
2005
2017
K
Endovascular aneurysm repair
(progression)
1997
2000
3,975
2017
L
Nociceptors/molecular pharmacology of opioid receptors
(progression)
1997
2001
1,812
2017
M
Protein engineering/two hybrid assay
(progression)
1997
2005
5,319
2017
N
Quality of life for HIV patients
(progression)
1997
2000
501
2017

**Fig. S11. Trajectories for setA breakthroughs**. Trajectories for setA breakthroughs described in table S2, excluding gold standard trajectories. **(A-N)** Nodes are shaded by % new publications and sizes are relative within each trajectory; the number of articles in the largest cluster for each trajectory is indicated; the number of articles in the largest cluster for each trajectory is indicated. Signal clusters are denoted with black asterisks and breakthrough clusters with light blue asterisks. Signal year, breakthrough year, and terminal year (2017) is indicated for each trajectory.

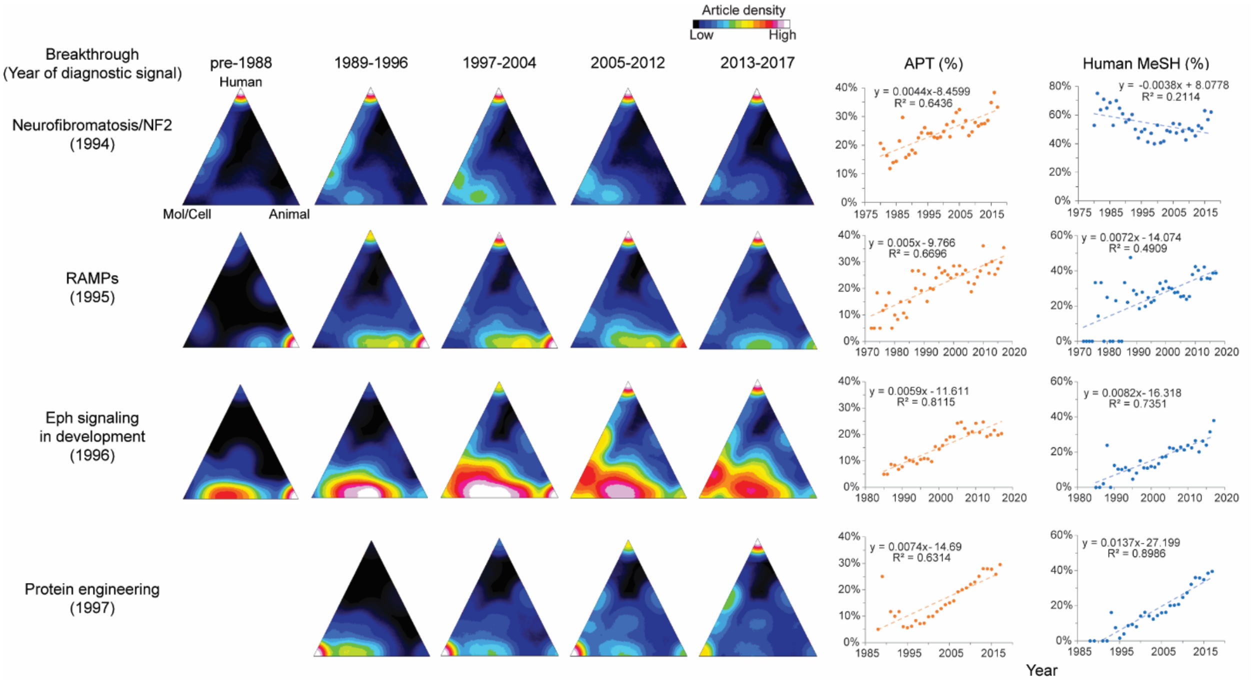


**Fig. S12. Translational progress for setA breakthroughs leading to drug development**. Snapshots of the triangle of biomedicine for publications in the citation networks of four setA breakthroughs that culminated in drug development (also see Movies S1-S4). Orientation of the Human, Animal, and Molecular/Cellular (Mol/Cell) vertices of the triangles is indicated on the left-most triangle for Neurofibromatosis/NF2. Growth in average Approximate Potential to Translate (APT)(*20*) and Human MeSH for each topic is also shown for each topic; trendline equations and $R^2$ values are displayed. As these areas of research make progress toward translation, the citation networks trend toward human-focused research, and average APT score increases.

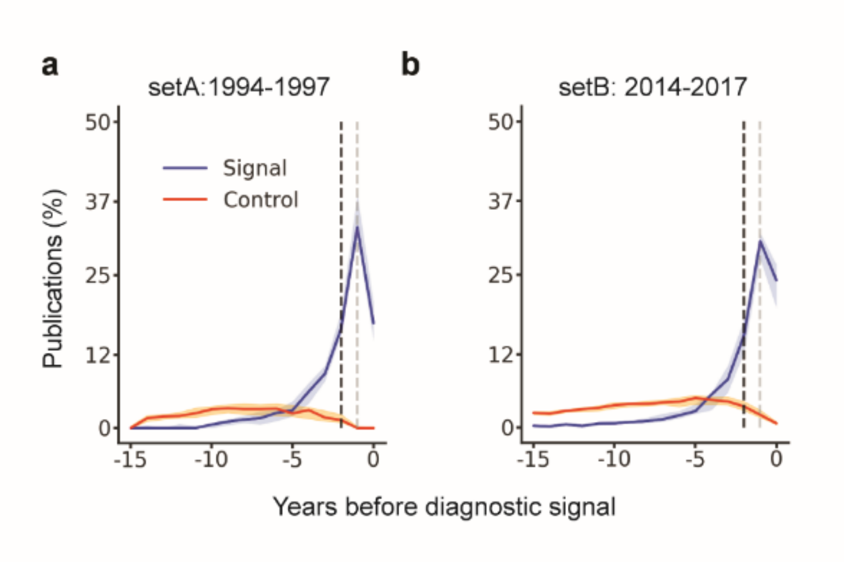


**Fig. S13. Dynamic behavior of regression features in clusters that signal future breakthroughs.** Comparison of (**A**) %New (percentage of all papers in a cluster that appeared no earlier than year x-1, where x is the terminal year of the cluster; grey dashed line) and (**B**) ancestral %New the percentage of all papers in year x-1 that appeared no earlier than year x-2 (black dashed line) for setA and setB clusters relative to their size-matched controls. For comparison of entropy and hRCR, see Fig. 4.

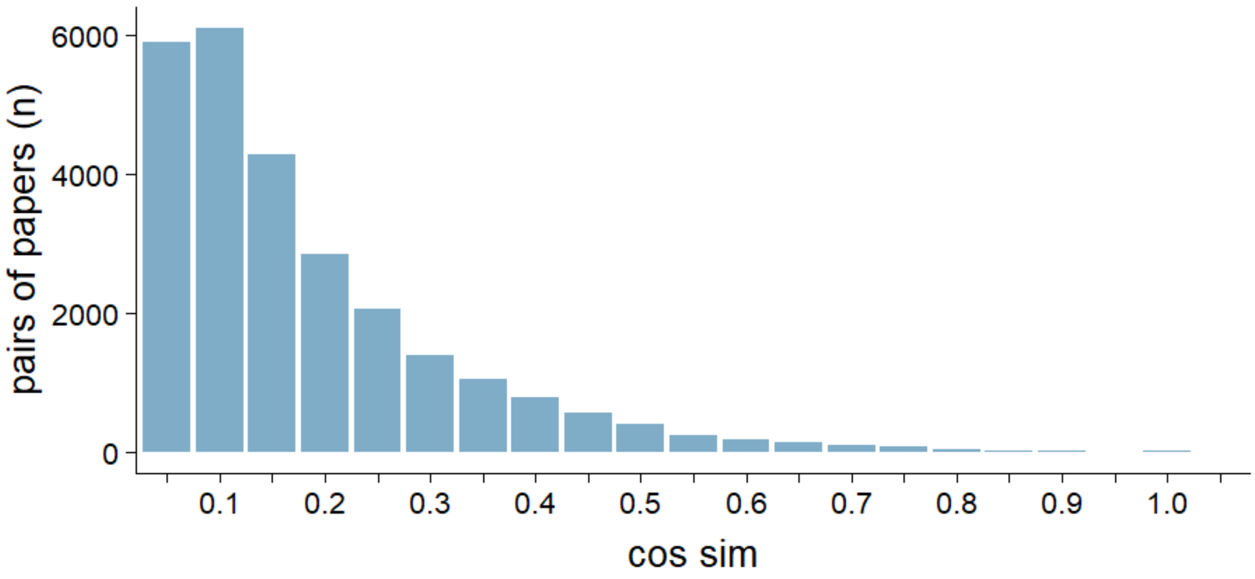


**Figure S14. Distribution of similarity for PMID pairs randomly sampled from the full CCN.** Values for cosine similarity (cos sim) are binned in increments of 0.05. No PMID appears more than once. The number of pairs at each similarity is shown (n = 26,330 pairs).

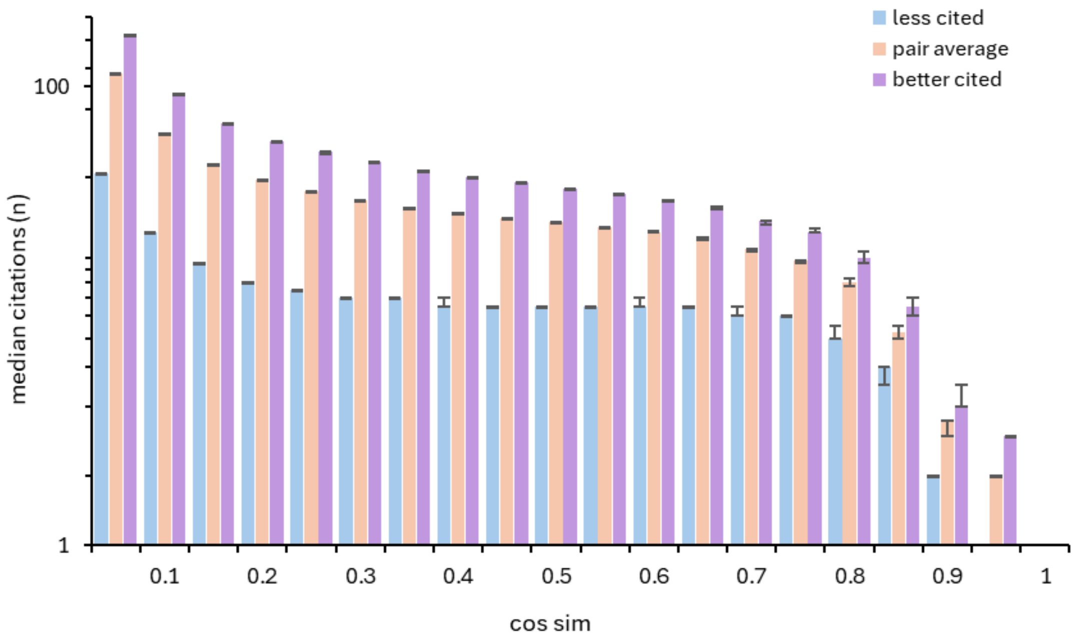


**Fig. S15. Semantic similarity between papers in a co-citation network as a function of citation counts.** 50,000 PMIDs that were cited at least once were randomly sampled from PubMed, and all of their partners in the full CCN were identified. Cosine similarity (cos sim) was calculated for every pair (approximately 37 million comparisons) and binned in increments of 0.05. Median citations in each bin are presented, considering either the number of citations received by the less-well cited PMID of the pair (blue bars), the average of both PMIDs in the pair (orange bars), or the better-cited PMID of the pair (purple bars). Error bars were computed by bootstrapping.

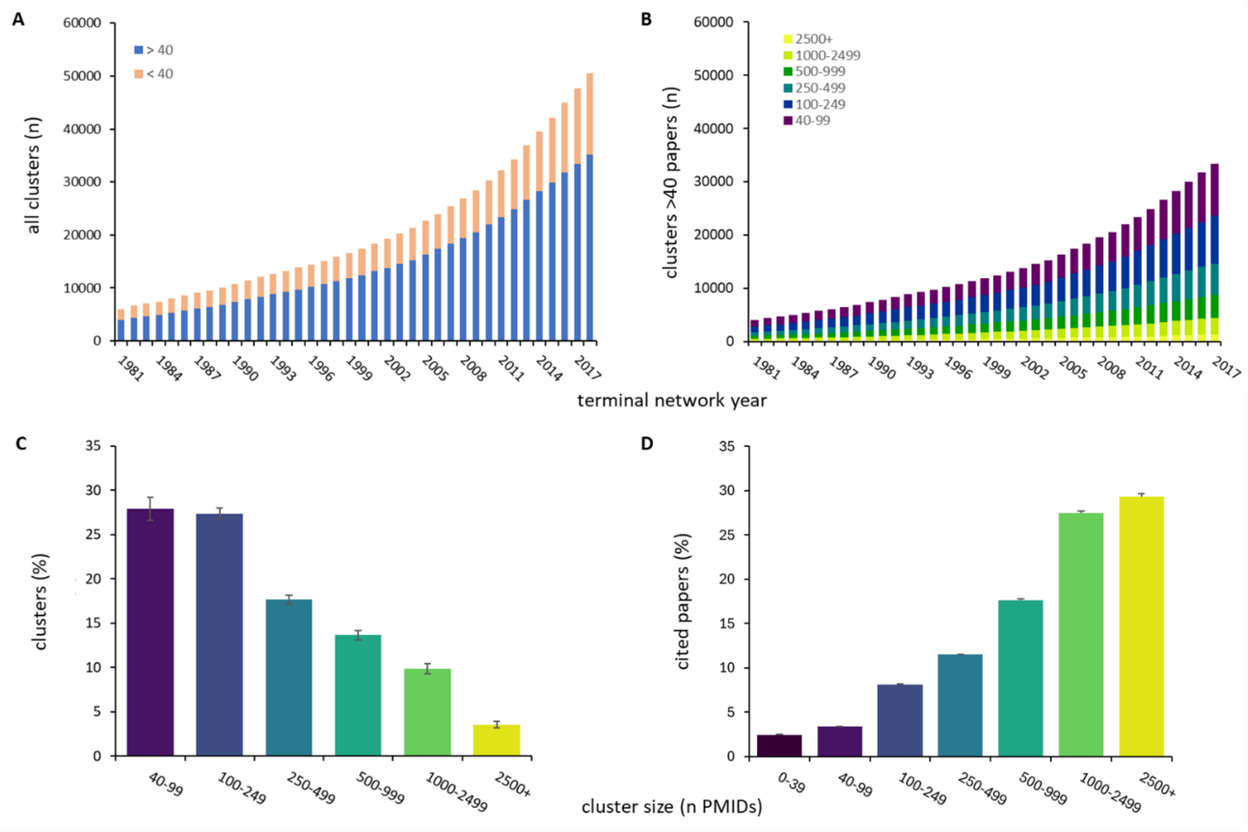


**Figure S16. Characteristics of RMCL/CCN clusters derived from historical networks.** For each historical network, **(A)** number of clusters (n) of < 40 or ≥ 40 papers, orange vs blue bars; **(B)** number of clusters (n) in each of six size categories; **(C)** percent of clusters in each size category, averaged over all network years; **(D)** percent of papers in each cluster size bin, averaged over all network years. Purple, 40-99 papers; blue, 100-249 papers; teal, 250-499 papers; green, 500-999 papers; light green, 1000-2499 papers; yellow-green, >2500 papers. Error bars, SEM.

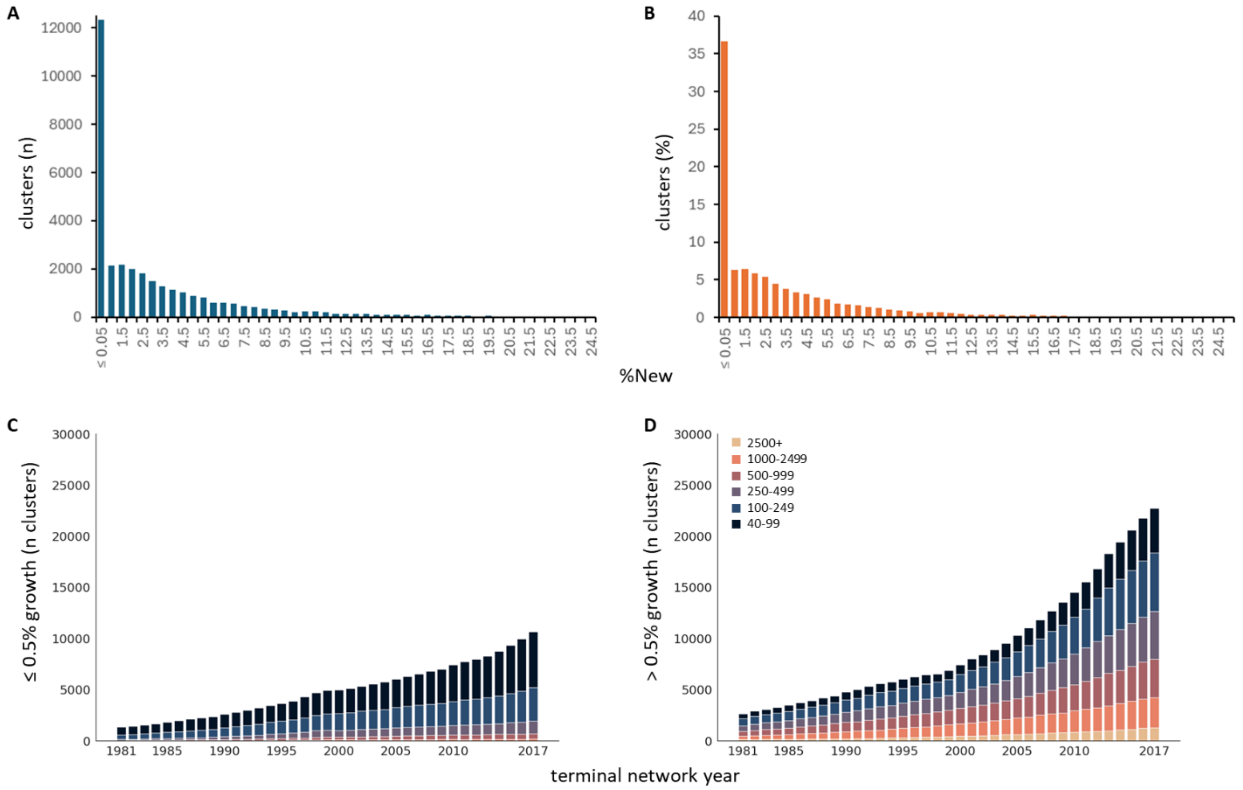


**Figure S17. Growth and stagnation of RMCL/CCN clusters.** (**A**, **B**) Number (blue bars) and percent (orange bars) of clusters at indicated values of %New in the 2017 network. Number of stagnant (**C**) and growing (**D**) clusters in each of six size categories (navy, 40-99 papers; blue, 100-249 papers; lavender, 250-499 papers; rose, 500-999 papers; orange, 1000-2499 papers; peach, >2500 papers) for each historical network.

| Gold standard[*] | Prize/Award | Trajectory type | Year of breakthrough[†] | Year of signal | Interval (years) | %New of signal[‡] |
|---|---|---|---|---|---|---|
| *Helicobacter pylori* | Phys. or Med. Nobel 2005 | divergence | 1984 | 1981 | 3 | 21.2% |
| Cystic fibrosis drugs | Alpert 2018 | convergence | 1990 | 1985 | 5 | 28.9% |
| Olfactory biology | Phys. or Med. Nobel 2004 | progression | 1991 | 1981 | 10 | 25.3% |
| Hypertension drug and nitric oxide | Phys. or Med. Nobel 1998 | progression | 1992 | 1985 | 7 | 37.5% |
| Vesicle trafficking | Phys. or Med. Nobel 2013 | convergence | 1993 | 1984 | 9 | 44.4% |
| Apoptosis/Bcl2/Bax | Wiley 2002 | convergence | 1993 | 1987 | 6 | 45.0% |
| Hepatitis C drugs | Phys. or Med. Nobel 2020 | convergence | 1993 | 1991 | 2 | 17.7% |
| Breast cancer genetics | Lasker 2014 | progression | 1995 | 1994 | 1 | 37.0% |
| Cancer immunotherapy | Phys. or Med. Nobel 2018 | progression | 1996 | 1993 | 3 | 32.8% |
| Carbon nanotubes | Physics Nobel 2010 | progression | 1997 | 1987 | 10 | 41.9% |
| RNA interference | Phys. or Med. Nobel 2006 | progression | 1998 | 1990 | 8 | 14.6% |
| TLR immunity | Phys. or Med. Nobel 2011 | divergence | 1998 | 1988 | 10 | 32.0% |
| Leptin/orexin | Lasker 2010 | convergence | 1998 | 1996 | 2 | 44.9% |
| Neurogenesis in adult humans | Keio Medical Science Prize 2008 | progression | 1998 | 1996 | 6 | 31.1% |
| EGFR drug targeting | Lasker 2019 | divergence | 2002 | 1990 | 12 | 35.3% |
| SRFM | Chemistry Nobel 2014 | progression | 2006 | 1996 | 10 | 53.1% |
| miRNA exosomes | ISEV Special Achievement Award 2015 | progression | 2007 | 2004 | 3 | 24.8% |
| iPSCs | Phys. or Med. Nobel 2012 | progression | 2008 | 2004 | 4 | 18.2% |
| Microbiome | Massry 2017 | convergence | 2008 | 2007 | 1 | 21.2% |
| Electronic delivery of mental health care/CBT | Nordic Medical 2014 | divergence | 2011 | 2010 | 1 | 20.1% |
| Gene editing | Chemistry Nobel 2020 | convergence | 2014 | 2008 | 6 | 32.5% |

**Table S1. Summary statistics describing gold standard breakthroughs.**

[*]Topics that have received formal recognition as breakthroughs by the scientific community.

[†]The table is organized in order of the year of the breakthrough.

[‡]The percentage of all papers in a cluster that appeared no earlier than year x-1, where x is the year of the cluster.

| Topic* | Prize/Award/<br>Drug development | Trajectory type | Year of breakthrough | Year of signal | Interval (years) | %New of signal | N pubs (signal cluster) |
|---|---|---|---|---|---|---|---|
| Breast cancer genetics | Lasker 2014 | progression | 1995 | 1994 | 1 | 37.0% | 474 |
| Human genome: cloning/sequencing methods | Gabbay Award 2000; Biotechnology Heritage Award 2001 | progression | 2001 | 1994 | 7 | 45.4% | 102 |
| Protein disulfide isomerase/ oxidative protein folding | AAAS/GE Young Life Scientist Award 2004 | progression | 1998 | 1994 | 4 | 60.6% | 92 |
| Neurofibromatosis/NF2 | Drug development | progression | 2013 | 1994 | 19 | 39.7% | 103 |
| Differential display (DD-PCR)/ plant evolution/ transgenic crops | Lavoisier Medal for Technical Achievement 2012 | progression | 2004 | 1994 | 10 | 43.8% | 320 |
| ALS/SOD | Breakthrough Prize in Life Sciences 2018 | progression | 2006 | 1995 | 11 | 51.5% | 158 |
| EuroSCORE | Pedro Cassio Foundation Award 2013 | progression | 1999 | 1995 | 4 | 42.7% | 126 |
| Adrenomedullin/RAMPs | Drug development | progression | 1998 | 1995 | 3 | 62.3% | 61 |
| Leptin/orexin | Lasker 2010 | convergence | 1998 | 1996 | 2 | 44.9% | 282 |
| SRFM/GFP | Chemistry Nobel 2008 | progression | 2006 | 1996 | 10 | 53.1% | 88 |
| PET-CT/efMRI | Killam Prize 2020 | progression | 1998 | 1996 | 2 | 29.7% | 353 |
| Receptor tyrosine kinases (RTKs) in development | Gruber Neuroscience Prize 2020 | progression | 1998 | 1996 | 2 | 37.3% | 283 |
| NMR analysis of protein dynamics | Laukien Prize 2004 | progression | 2005 | 1996 | 9 | 38.3% | 131 |
| Endovascular aneurysm repair | Jacobsen Innovation Award 2017 | progression | 2000 | 1997 | 3 | 26.4% | 278 |
| Nociceptors/molecular pharmacology of opioid receptors | Julius Axelrod Award 2012; Gruber Neuroscience Prize 2023; National Medal of Science 2023 | progression | 2001 | 1997 | 4 | 65.6% | 148 |
| Protein engineering/two hybrid assay | Chemistry Nobel 2018 | progression | 2005 | 1997 | 8 | 55.5% | 152 |
| Quality of life for HIV patients | NA | progression | 2000† | 1997 | 3 | 44.3 | 97 |

**Table S2. Summary statistics describing setA breakthroughs.**

*Topics predicted to be breakthroughs by diagnostic signals (1994-1997), excluding a false positive signal (3rd gen coxibs). Topics in the grey rows also appear in the collection of gold standards.

†Hays *et al*. report the first large-scale evaluation of the health-related quality of life of HIV-positive adults relative to the general US population or patients with other chronic conditions (2000 Am. J. Med. 108(9):714-22).

| Topic* | Prize/Award | Year of breakthrough | Year of signal† | Interval (years) | %New of signal | N pubs (signal cluster) |
|---|---|---|---|---|---|---|
| Histone modifications | Lasker 2018 | 1988 | 1982 | 6 | 41.0% | 76 |
| DNA damage response | Lasker 2015 | 1993 | 1984 | 7 | 33.6% | 742 |
| Autophagy | Medicine 2016 | 1992 | 1984 | 8 | 44.3% | 563 |
| AlphaFold | Lasker 2023 | 2010 | 1987 | 23 | 44.6% | 200 |
| Mechanoreceptors & thermoreceptors | Medicine 2021 | 1997 | 1989 | 8 | 100% | 160 |
| TOR kinase | Lasker 2017 | 1996 | 1992 | 4 | 35.4% | 331 |
| HIF1 | Lasker 2016; Medicine Nobel 2019 | 1991 | 1992 | <1‡ | 35.9% | 788 |
| mRNA vaccines | Medicine 2023 | 2005 | 1992 | 13 | 34.3% | 424 |
| Unfolded protein response | Lasker 2014 | 1993 | 1994 | <1‡ | 36.3% | 116 |
| Optogenetics | Lasker 2021 | 2003 | 1996§ | 7 | 53.1% | 88 |
| Neurology of spatial positioning | Medicine 2014 | 1971 | NA | NA | NA | NA |
| aDNA | Medicine 2022 | 1997 | 1998 | <1‡ | 45.8% | 28 |

**Table S3. Summary statistics describing recognized breakthroughs not used as gold standards.**

*Topics that would have been predicted in advance by our approach that subsequently went on to receive formal recognition as breakthroughs by the scientific community.

†The table is organized in order of the year of the signal. Topics in the grey rows scored positively in our logistic regression but were below our size threshold and/or were slow growing.

‡Relevant breakthrough publications appeared in the second half of the year and were rapidly cited, resulting in a near-simultaneous signal.

§Branches off SRFM/GFP trajectory in 2005.

**Movies S1-4. Translational progress for setA breakthroughs leading to drug development**. Timelapse of the triangle of biomedicine for publications in the citation networks of four setA breakthroughs that culminated in drug development: Neurofibromatosis/Nf2 (Movie S1); RAMPs (Movie S2); Eph signaling in development (Movie S3); Protein engineering (Movie S4). Movies correspond to still images from fig. S8.

**Data S1. Agreement between subject matter experts and RMCL cluster designations.** Subject matter experts (SMEs) were asked to partition publications assigned to pairs of RMCL clusters with increasing word2vec similarity. Results are presented in order of topics determined by word2vec to be least similar (similarity 0.0) to most similar (similarity 0.9).

**Data S2. Full network and RMCL statistics (1981-2017).** Summary statistics for articles published between 1981 and the reference year.

**Data S3. Breakthrough publications for the SRFM trajectory.** The 279 publications in the SRFM trajectory authored by the Nobelists credited with the transformative breakthrough.

**Data S4. Features of gold standards, predict breakthroughs, and clusters that lack a breakthrough signal.** Values for all metrics used to select breakthrough predictions, including the four components of the logistic regression signal, %NIH funding, and cluster size. Values for gold standards and for a random selection of clusters that were not identified by the logistic regression are listed for comparison.

**Data S5 and S6. Publications in gold standard and setA signal clusters with a rapid growth in influence.** List of publications that at least tripled in historical RCR from two years prior to the signal cluster and had an RCR of at least 3.0 by the signal year for gold standards (Data S4) and setA (Data S5).

**Data S7. Glossary of all features tested for statistically significant differences in the logistic regression approach.**

**Supplementary Text**

**Sample calculation of CCN cosine similarity**

The table below shows hypothetical corpus composed of a set of eight papers (111, 123, 400, 454, 500, 555, 600, 700), each of which cites three to six papers:

Paper 111 → reference list: [100, 99, 98, 94]

Paper 123 → reference list: [94, 33, 22, 11]

Paper 400 → reference list: [94, 33, 11]

Paper 454 → reference list: [100, 22, 10]

Paper 500 → reference list: [400, 123, 33, 17]

Paper 555 → reference list: [98, 95, 94, 22, 11, 10]

Paper 600 → reference list: [94, 33, 11]

Paper 700 → reference list: [77, 44, 18, 17, 11]

Paper 11 is co-cited with paper 10 once; with paper 17 once; with 18 once; with 22 twice; with 33, 3 times; with 44 once; with 94, 4 times; with 95 once; and with 98 once. For [p,n] where p is the paper co-cited and n is the number of times it is co-cited, and including the number of times the paper itself is cited, the sparse vectors that describes papers 11 and 22 would be represented as

11 → [10,1], [17, 1], [18, 1], [22, 2], [33,3], [44, 1], [94, 4], [95, 1], [98,1], [11,5]

22 → [10,2], [11,2], [33,1], [94,2], [95,1], [98,1], [100,1], [22,3]

Calculation of the CCN cosine similarity for papers 11 and 22 would be as shown below:

$$cosineSim(ccn_{11}, ccn_{22}) = \frac{ccn_{11} \bullet ccn_{22}}{|ccn_{11}||ccn_{22}|}$$

The numerator is the dot product (that is, the sum of the products) of non-zero elements (defined as those elements that exist in both vectors) and the denominator is the square root of the sum of each element squared; for the values given, the solution of the above equation is approximately 0.8. As citation of a paper increases, the magnitude of the vector will increase as well, so that a paper with more citations will be normalized by the larger size of the denominator. This method also has the added advantage of improving the temporal cohesiveness of linked articles, relative to either a simple CCN or a direct citation network (fig. S1).

**Selection of thresholds for generation of co-citation networks and RMCL clusters**

CCNs are very dense, i.e., the nodes in a CCN are connected by a large number of edges. For the full CCN of all 18.5 million papers deposited in PubMed through the end of 2017, the median node degree is approximately 17,000. The discovery and analysis of community structure in such a large, dense network is difficult, in part because the amount of time required to calculate all possible ways to divide the graph increases exponentially with the size of the network. However, not all edges in a CCN are equal in terms of their informational content. We therefore reasoned that it should be possible to reduce the number of edges to a computationally tractable number, while retaining the most significant information in the network, i.e., those edges that connect papers with the greatest degree of similarity.

Where paper R cites papers (A,B) and paper Q cites papers (B,C), the full CCN for any one of the three cited papers is (A,B,C; fig. S2A). However, A and C do not appear together in the reference list of either R or Q; their connection is transitive. In contrast, the pairs (A, B) and (B,C) do appear alongside each other in a single reference list and may be considered connected by first-order edges. Limiting to first order edges, which presumably have the greatest degree of similarity and therefore the most information about community structure, reduces the median node degree of the graph of PubMed to approximately 250. Each paper in the full graph can be represented as a sparse vector that contains an element for each of its co-cited papers; the value of those elements equals the number of times a given paper is co-cited with the paper described by the vector, as outlined in the above example. A similarity score can then be calculated between any two papers.

Euclidean distance is sensitive to vector magnitude and therefore not a good candidate for our use case. We also ruled out overlap, also known as Szymkiewicz–Simpson, coefficients as unsuitable, since this measure would introduce distortions from small n effects when poorly cited papers were considered in the denominator. Jaccard similarity also generates a skewed measurement when the denominator or numerator are of very different sizes. We therefore chose cosine similarity as the most logical approach, although extensive testing of alternative measures might result in further optimizations. Of all edges with a cosine similarity of ≥ 0.35 in the full, un-pruned CCN of PubMed, 93% are first order edges (fig. S2B). Applying a cosine similarity cutoff of 0.35 to first order edges in the network brings the median node degree to 20; the RMCL algorithm operates best on networks with a median node degree ≤ 100 (*92*).

The inflation parameter of the RMCL algorithm controls the extent to which weak connections in the network are preserved in the subgraphs; the lower the inflation, the more edges are retained. Inflation settings in the range of 1.2-5.0 are typically advised (*92*). We reasoned that the more we prioritized the maintenance of weak connections, the less likely we would be to over-split the graph, i.e., the less likely we would be to assign papers that SMEs would identify as one topic to two clusters. However, the greater the number of weak connections that are retained, the more papers will be trapped in extremely large clusters that are unlikely to accurately represent a single field. At an inflation setting of 1.1, across all network years over 80% of cited papers (~15 million) are trapped in a single large cluster; at an inflation setting of 1.2, less than 3% are trapped (fig. S3). While further refinements might be realized by an in-depth analysis of inflation settings between 1.1 and 1.2, such a comprehensive study carried out at scale would be beyond the scope of any single manuscript. We therefore chose to proceed with an inflation setting of 1.2, since increasing to 1.3 would increase the risk of over-splitting small topics, which we theorized were more likely to represent promising emerging areas of research.

It should be noted that, as is true of conventional CCNs, our method generates a network where highly cited papers are overall less similar to their co-cited partners than less well-cited papers are to theirs (fig. S15). We would expect those lower-similarity edges to exist at a lower density and correspond to longer path lengths that are targeted by RMCL, so that influential papers operate like high-mass objects in a gravitational field, nucleating related work and existing relatively rarely in small clusters or in isolation. To test this hypothesis, we randomly selected 200,000 papers, half each from one of two groups – papers ranking in the top quintile of RCR values (3.0 or greater), and papers ranking around the $50^{th}$ percentile (RCR between 1.05 and 0.95). As expected, we found that influential papers are more likely to be assigned to clusters that meet our threshold for a cohesive topic (40 or more papers; 96,204 vs. 95,541, $p = 9.9 \times 10^{-14}$, chi-square with Yates continuity correction). At the same time, poorly cited papers (RCR 0.05-0.25) are more likely to be found outside a coherent topic cluster (89,603 vs. 95,541, $p < 2.2e-16$, chi-square with Yates continuity correction) in spite of their higher similarity to co-cited works. In this way, differences in similarity between CCN partners across the influence spectrum helps to generate topics that mirror those conceived of by human experts.

**Characteristics of the corpus**

At an inflation setting of 1.2, RMCL processing of the 19.3M cited papers in PubMed, beginning with the oldest publications in the database and running through the end of 2017, generates 33,392 CCN clusters of more than 40 papers. 23,867 of these (more than 70%) are in the largest connected component of the network (Fig. 1D);

89% (17.2M) of all cited papers are distributed over the clusters that make up the LCC. Although it represents a large majority of all clusters, it should be noted that a cluster does not need to be part of the LCC in order to part of a historical trajectory, and that our regression analysis considers all clusters of more than 40 papers, regardless of their membership in the LCC. Over the time period of our analysis the number of clusters in PubMed has grown exponentially, increasing by approximately 10-fold (fig. S16A,B). However, the distribution of clusters of all sizes remains remarkably stable (fig. S16B,C). Across all 37 historical CCN/RMCL networks, each of which excludes all papers and citations beyond the terminal year (Data S2), slightly less than a third (30 ± 0.3%) of clusters are small (>40 papers; fig. S16A). These small clusters contain very few papers; 94±0.5% of cited papers are assigned to a cluster that meets our size threshold (> 40; fig. S16D).

12,334 clusters in the 2017 network experienced very low or no growth, i.e., added few or no new papers (≤0.5%New) in their terminal year (fig. S17A). In the 2017 corpus, these stagnant clusters contain 11.9% of all cited papers, whereas less than 5% of clusters of >40 papers had a %New value of 14.5% or greater (fig. S17B). For comparison, the range of %New in our 21 gold standard signal clusters was 14.5-52.6%. Perhaps unsurprisingly, across all network years stagnant clusters tend to be smaller than those that are actively growing (fig. S17C, D).

**Stability of clusters and trajectories over time**

Any given scientific topic relies on a core collection of scholarly literature that practitioners continue to recognize as relevant over time. Thus, to accurately represent reality, the agnostic computational identification of distinct scientific topics from an unstructured database such as PubMed must produce groups of papers that stay together even as new literature is added to the full corpus each year. Having chosen CCNs as the most advantageous method for representing connections between papers (see Main Text for details), we asked which of two approaches generates more stable clusters: our implementation of RMCL (see above and Main Text) or the well-recognized Leiden algorithm (*93*), which uses a combination of local and random movement of vertices between sub-graphs to identify sub-communities within large, complex networks.

Using the Leiden algorithm with a low-resolution setting of 0.0002 to minimize the fragmentation of the whole CCN of PubMed, generates clusters in year x that can be mapped to a direct descendant in year x+1 an average of 80% of the time, where a direct descendant is defined as one that shares 50% or more of its papers (sum of blue and orange bars, fig. S4B). By the same measure, more than 95% of RMCL-generated clusters can be mapped to a direct descendant at an inflation setting of 1.2 (sum of blue and orange bars, fig. S4A). The fraction of direct descendants that capture the entirety of its chronological predecessor (orange bars, Figure S4) is approximately equivalent for the two methods (19% for the Leiden algorithm vs 16% for RMCL, on average over all years), while the fraction that split, shed more than half of their papers, or disintegrate completely from one year to the next is more than 6x higher for the Leiden algorithm (on average 22% vs. 3.2% for RMCL). Taken together, these data show that RMCL outperforms the Leiden algorithm in its ability to produce clusters that were stable from one year to the next.

We next asked how stably papers are assigned to our RMCL-defined clusters over multiple years. We began with papers published in 1981, then selected a partner paper assigned to the same cluster in our 1984 network. Since citations take time to accrue, peaking approximately three years after publication in biomedical disciplines (*94*), this is the earliest point from which we can accurately trace the fraction of pairs that stay in the same cluster over time. We found that papers initially assigned to the same cluster have a strong tendency to stay together; 80% still share a cluster after more than thirty years. Even roughly half of the most semantically distant pairs selected at random still share a cluster after thirty years (fig. S6A).

Together our data suggest that once formed, individual RMCL/CCN clusters are typically quite stable (fig. S4, fig. S6A). However, from one year to the next a small percentage of clusters lose more papers than they gain. This may happen when new subfields begin to evolve from a common ancestor, or as in the case of SRFM/GFP (Fig. 1E), when subgroups of investigators in a single field change their focus from one aspect of a problem to another, resulting in a detectable divergence in co-citation patterns. Our design choice to match clusters across years by a single, simple rule – that they must share at least 50% of their papers – means that the size of a descendant cluster could theoretically decrease by up to 49%, relative to its historical predecessor. Large changes in size are uncommon; splits in the 45-55% range account for 2.5% of all cases, while less than 0.08% of clusters in year x+1 arise from a 50:50 partition of a cluster in year x. Although growing clusters are more likely to exhibit a change in size, the size of stagnant clusters may decrease if scientists reinterpret or discover new relevance for a subset of their papers. In this way, the growth and change of CCNs over time accurately captures the dynamic nature of science (*31*).

Having quantified the stability of individual clusters, we next asked about the stability of entire trajectories. To answer this question, we built trajectories for all clusters in the 2017 historical network, extending back to the birth of the topic they represent. We could then ask what fraction of each individual publication's lifetime was spent in the trajectory terminating in its assigned cluster in 2017. Short trajectories are more common than long ones; approximately 10% of papers are assigned to a trajectory with a lifetime of only two or three years. This generates relatively small but detectable groups of publications that spend 50% of their lifetime in a two-year, or 33% and 67% in a three-year, trajectory (fig. S6B). A much larger group of papers (54%) remain in a single trajectory for upwards of 80% of their lifetime; almost a third (30%) never leave the trajectory to which they are initially assigned (fig. S6B). Overall, our data suggest that clusters and trajectories are stable enough to accurately describe existing scientific topics and dynamic enough to capture the evolution of new ones.
