## Supplementary figures and images for "Prediction of transformative breakthroughs in biomedical research"

### Supplemental Movie 1

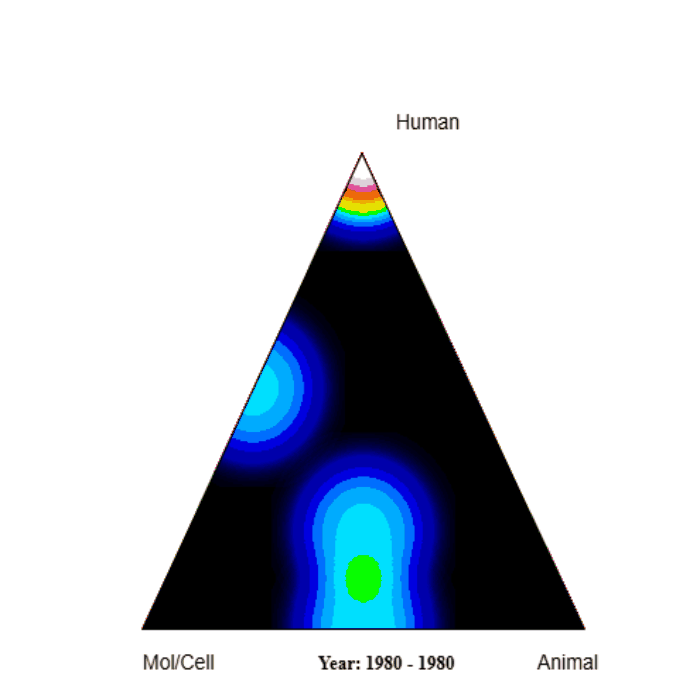

### Supplemental Movie 2

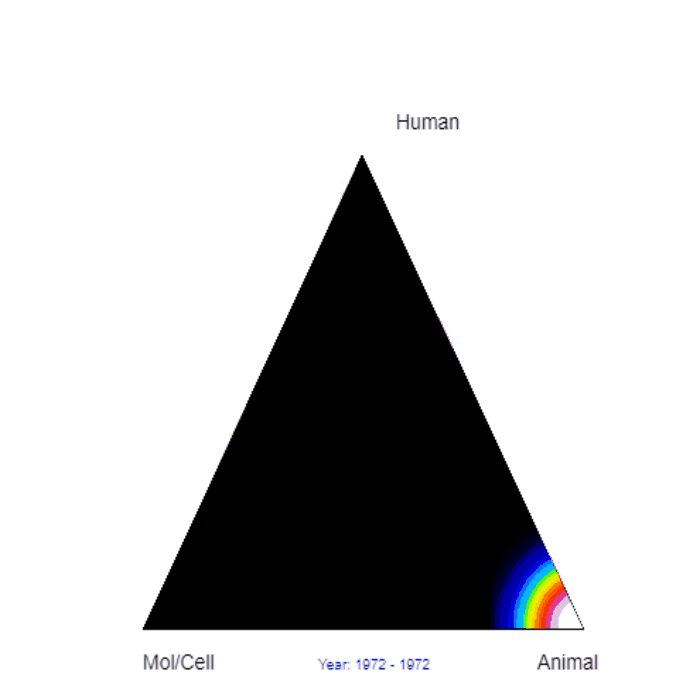

### Supplemental Movie 3

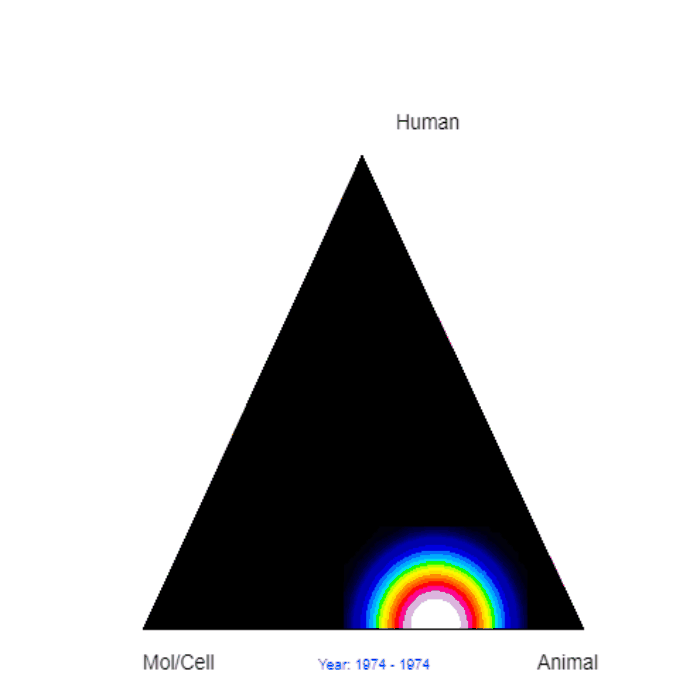

### Supplemental Movie 4

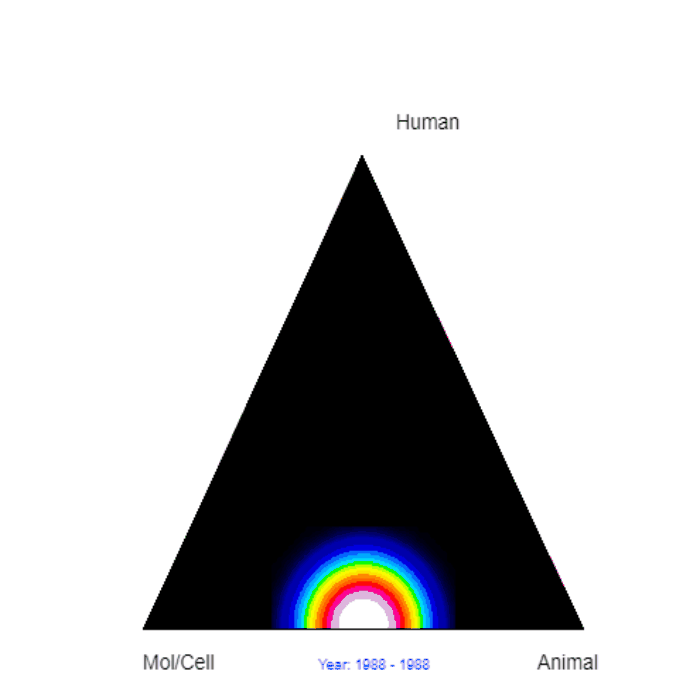
